# Three-dimensional fossils of a Cretaceous collared carpet shark (Parascylliidae, Orectolobiformes) shed light on skeletal evolution in galeomorphs

**DOI:** 10.1101/2025.03.14.643294

**Authors:** R.P. Dearden, Z. Johanson, H.L. O’Neill, K. Miles, E.L. Bernard, B. Clark, C. Underwood, M. Rücklin

## Abstract

A rich fossil record of teeth shows that many living shark families’ origins lie deep in the Mesozoic. Skeletal fossils of the sharks to whom these teeth belonged are far rarer and when they are preserved are often flattened, hindering understanding of the evolutionary radiation of living shark groups. Here we use computed tomography to describe two articulated Upper Cretaceous shark skeletons from the Chalk of the UK preserving three-dimensional neurocrania, visceral cartilages, pectoral skeletons and vertebrae. These fossils display skeletal anatomies characteristic of the Parascylliidae, a family of Orectolobiformes now endemic to Australia and the Indo Pacific. However, they differ in having a more heavily mineralised braincase and a tri-basal pectoral fin endoskeleton, while their teeth can be attributed to a new species of the problematic taxon *Pararhincodon*. Phylogenetic analysis of these new fossils confirms that *Pararhincodon* is a stem-group parascylliid, providing insight into the evolution of parascylliids’ distinctive anatomy during the late Mesozoic-Cenozoic shift in orectolobiform biodiversity from the Northern Atlantic to the Indo Pacific. Meanwhile both *Pararhincodon* and extant parascylliids have a distinctive vertebral morphology previously described only in Carcharhiniformes, contributing a skeletal perspective to the picture emerging from macroevolutionary analyses of coastal, small-bodied origins for galeomorphs.

## Introduction

The evolutionary history of sharks (Selachii) is an emerging focus of macroevolutionary studies (Marion et al., 2024; Sorenson et al., 2014; Sternes et al., 2024) but is obscured by a sparse record of skeletal fossils. Because of their high evolutionary distinctiveness (Stein et al., 2018) and diversity in form and habitat yet tractable size (López-Romero et al., 2023), sharks are increasingly used as a group in which to understand the interplay of phenotype, habitat, and diversity through time (Marion et al., 2024; Sorenson et al., 2014; Sternes et al., 2024), and are moreover important from an ecological and conservation perspective (Dedman et al., 2024). Underlying these studies is a rich fossil record of isolated teeth, which shows that the selachian crown-group stretches far back into the Mesozoic (Underwood, 2006) and are used to constrain the evolutionary history of the group in time. However, there is a limit to the information that teeth and their limited character sets can provide for estimates of phylogeny and evolutionary timing (Brée et al., 2022), and the cartilaginous skeletons of these early crown-group selachians, which would provide more information, have a low preservational potential. Even in those Mesozoic Fossil-Lagerstätten that do preserve potentially information-rich skeletons, these fossils are often flattened (George et al., 2024; Shimada, 1997; Shimada & Cicimurri, 2005; Villalobos-Segura et al., 2023; Vullo et al., 2024), making it difficult to extract the level of detailed three-dimensional (3D) skeletal data that has demystified much of the record of sharks’ older, Palaeozoic relatives (Coates et al., 2017; Dearden et al., 2024; Pradel et al., 2014). This restricts understanding of key phylogenetic problems at the roots of selachians (Villalobos-Segura et al., 2022), limits the ability to incorporate traits into macroevolutionary studies (Marion et al., 2024) and obscures the morphological evolution that led to modern shark biodiversity.

The Upper Cretaceous Chalk of the UK is one of few locations to preserve numerous crown-group selachian cartilaginous skeletons three-dimensionally (Guinot et al., 2013; Maisey, 1985; Woodward, 1911), yet their skeletal anatomy has received little attention. The Chalk of the UK represents a shallow, subtropical ocean setting, and has been recognised as a Fossil-Lagerstätte for bony fishes (Friedman et al., 2015). Although it preserves a number of crown-group selachians (Guinot et al., 2013; Woodward, 1911), detailed description of their skeletal morphology has been limited to the problematic taxon *Synechodus* (Maisey, 1985, 2008), a member of the now-extinct Synechodontiformes (Klug, 2010). As a result, the skeletal anatomy of almost all of the Chalk selachians remains poorly understood compared to detailed descriptive work on flattened Upper Cretaceous elasmobranchs from Lebanon (George et al., 2024; Pfeil, 2021) and three-dimensional Upper Cretaceous batoids from Moroccan chalk (Claeson et al., 2013; Villalobos-Segura et al., 2019). Moreover the 3D imaging methods that have led to an the increasingly sophisticated understanding of extant elasmobranchs skeletal morphology (Dearden et al., 2021; Denton et al., 2018; Maisey, 2004; Staggl et al., 2022), and their Palaeozoic forebears (Coates et al., 2018, 2019; Dearden et al., 2024; Frey et al., 2019; Maisey, 2011) have yet to be deployed on these Chalk taxa. The Chalk provides a unique opportunity to characterise the skeletal anatomy of extinct crown-group selachians and help understand the complex evolutionary history of their living relatives (Boyd & Seitz, 2021).

Although the fishes of the Chalk are preserved three-dimensionally they are difficult to study using conventional means (Friedman et al., 2015). Here we use computed tomography to describe the skeletal anatomy of two sharks from the Chalk. We use skeletal characters to place these in a selachian phylogeny and show that these belong to a new species of stem-group parascylliid, sister group to other Orectolobiformes. We combine this with a new computed tomographic dataset for living parascylliids to show that these extinct parascylliids preserve plesiomorphic characteristics lost in living parascylliids, with implications for the early evolution of galeomorphs.

## Materials & Methods

### Institutional Abbreviations

CSIRO: CSIRO, Australian National Fish Collection, Hobart, Australia MNHN: Muséum national d’histoire naturelle, Paris, France

NHMUK: Natural History Museum, London, United Kingdom

### Fossil Specimens

Two principal fossil specimens are described here, both held at the Natural History Museum, London (Figure 1, Supplementary Figures 1, 2). At the time of writing NHMUK PV P 17223 is recorded as “*Synechodus*” (Figure 1a). It comprises a main block containing most of the fossil (NHMUK PV P 17223), an isolated tooth extracted from the matrix (NHMUK PV P 17223a), and a dermal denticle extracted from the matrix (NHMUK PV P 17223b), as well as several fragments broken from the main block which additional scanning shows contain dermal denticles and cartilage fragments. It is listed as being collected from the *Actinocamax quadratus Z*one (Jukes-Browne & Hill, 1904) of Britford, near Salisbury, by Dr. H.P. Blackmore. The *Actinocamax quadratus* Zone corresponds to the modern *Offaster pilula* and *Gonioteuthis quadrata* zones (Mortimore, 1986), which in turn correspond to the early Campanian (Guinot et al., 2013). NHMUK PV P 73821 (Figure 1b) is recorded as “Chondrichthyes”, and comprises a main block containing most of the fossil (NHMUK PV P 73821a), two samples of dermal denticles (NHMUK PV P 73821b+c), and an isolated tooth extracted from the matrix (NHMUK PV P 73821d). The only information of its provenance is that it was collected in Newhaven, which would be consistent with a Campanian or Santonian age (Mortimore, 2011). This should be confirmed more precisely in future through biostratigraphic correlation of nannofossils (Bramlette & Riedel, 1954).

**Figure 1.**
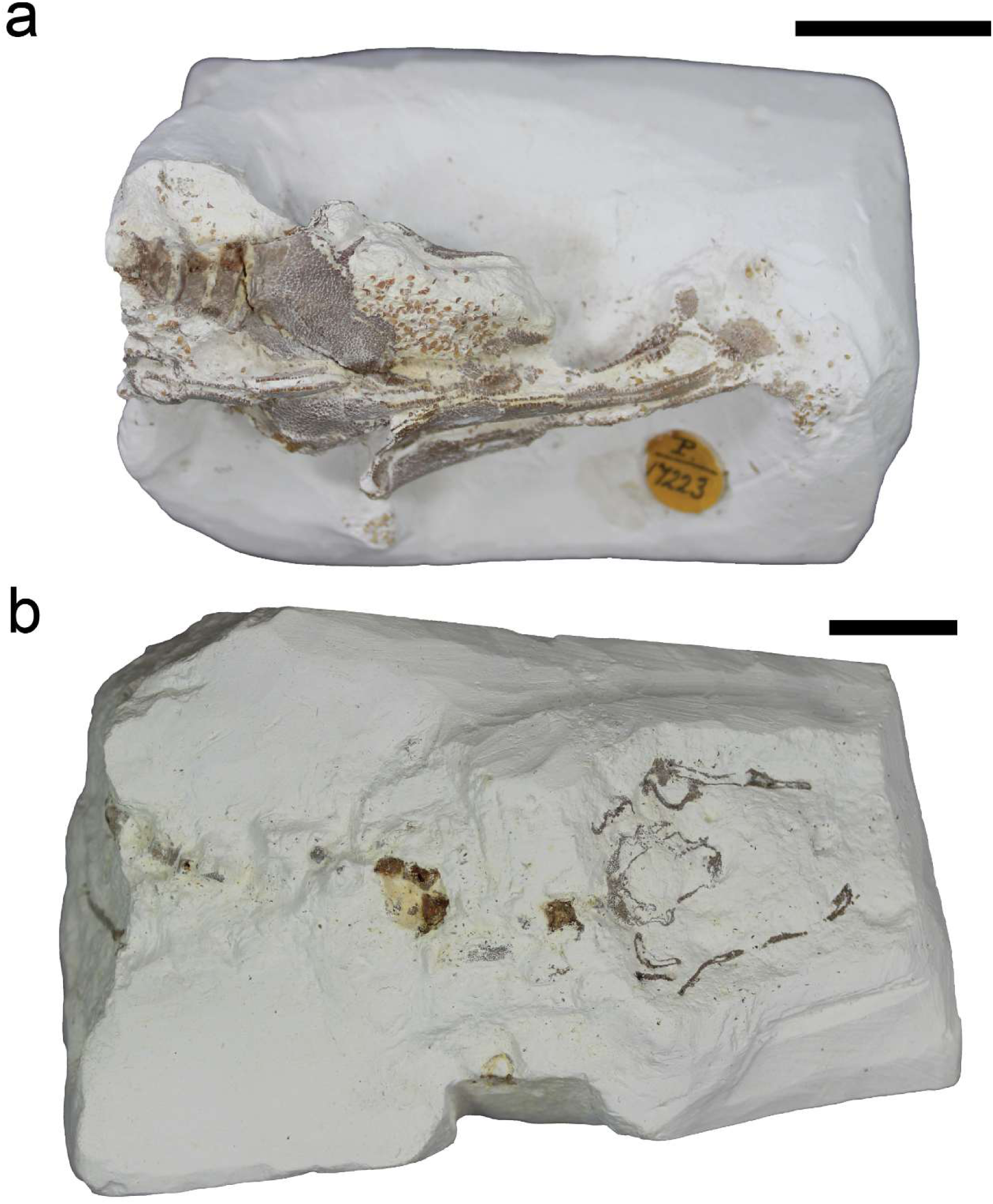
Photographs of the two fossils described in this study. a, NHM PV P 17723; b, NHM PV P 73821a. Photographs taken before removal of isolated teeth from matrix. Scale bars 10 mm.

### Extant *Parascyllium*

To aid interpretation of the fossils we studied the skeleton of an extant parascylliid shark *Parascyllium variolatum* CSIRO CA 3311 (Supplementary Figure 3). CSIRO CA 3311 is a 627 mm TL adult male collected onboard the Fishery Research Vessel *Solea* by demersal trawl on the 3rd Dec 1981 off southern Australia, 32°58’12.0”S 128°59’47.8”E at 72 metres depth.

### Micro-CT (computed tomography) scanning

Fossil specimens were scanned at the Natural History Museum, London, on a Nikon XT H 225 ST CT scanner. NHMUK PV P 17223a was scanned at 210 kV and 524 μA at a voxel size of 29.99 μm. A tungsten rotating reflection target was used with a 1 mm Sn filter, a 250 ms exposure and 3409 projections. NHMUK PV P 73821a was scanned in two sections, each at 155 kV and 129 μA at a voxel size of 31.72 μm. A tungsten reflection target was used with a 0.5 mm Al filter, a 500 ms exposure and 3700 projections. Extant specimen CSIRO CA 3311 was scanned at The University of Queensland, Australia by the Centre for Advanced Imaging on a Yxlon-Comet FF35. Two different scans were carried out: the first targeted the whole body, with 86 kV, 370 μA, 200 ms exposure time, and a 0.5 mm Al filter achieving a voxel size of 85 μm; the second targeted the head and pharynx, with 80 kV, 370 μA, 333 ms exposure time, and a 0.5 mm Al filter achieving a voxel size of 25 μm. Scan data were segmented in Materialise Mimics v25.0 (Materialise Software, Leuven, Belgium, https://www.materialise.com/en/healthcare/mimics), using manual thresholding as well as interpolation. All images of 3D data (Figure 2) were rendered in Blender v4.2 (blender.org).

**Figure 2.**
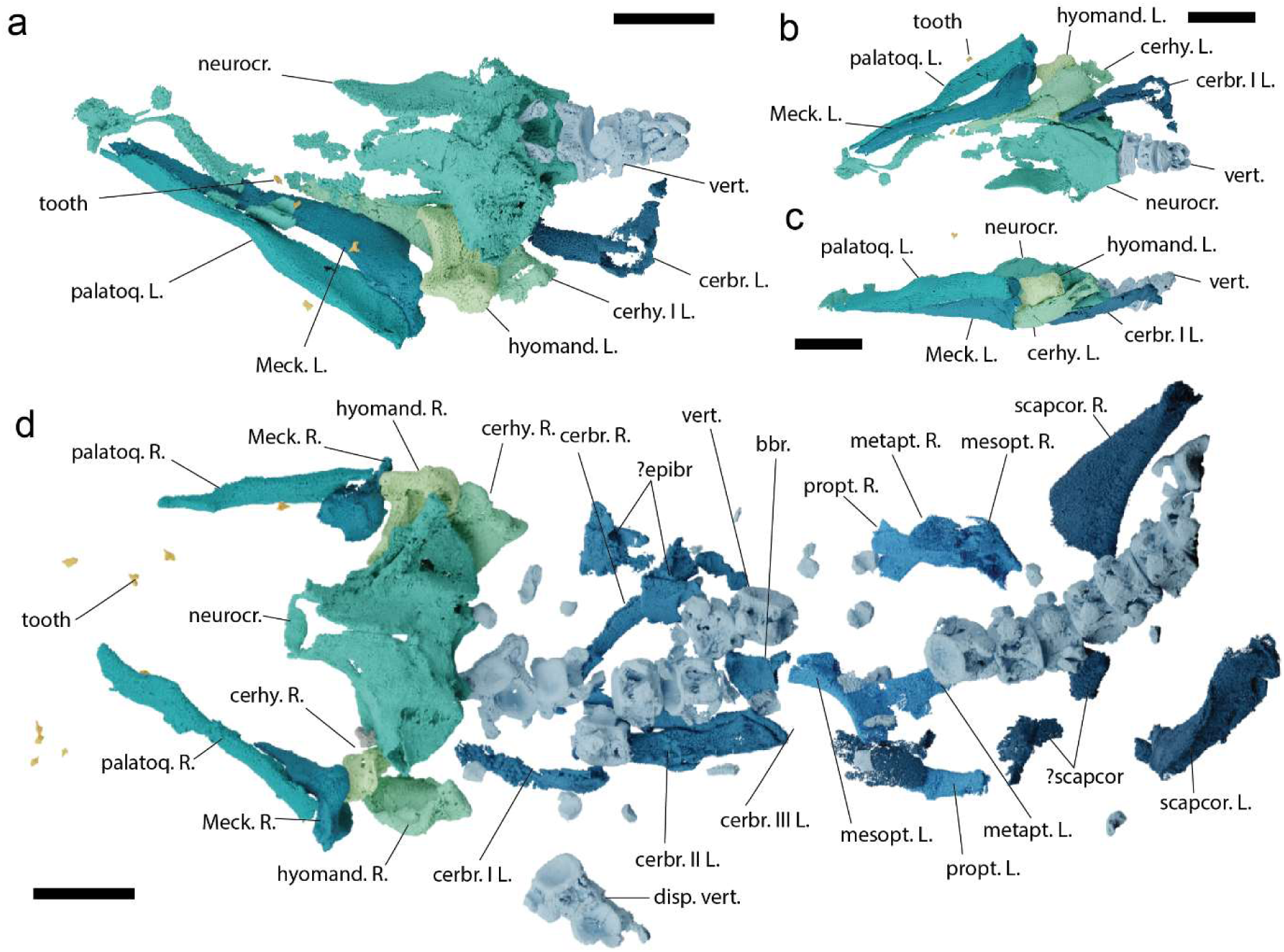
The skeletal anatomy of *Pararhincodon n. sp.* based on computed tomography. a-c The head of NHMUK PV P 17723 in dorsal (a), ventral (b), and left lateral (c) views; d, The head and thorax of NHMUK PV P 73821a in dorsal view. Abbreviations: bbr., basibranchial; cerbr., ceratobranchial; cerhy., ceratohyal; disp., displaced vertebrae; epibr., epibranchial; hyomand., hyomandibula; L., left; Meck, Meckel’s cartilage; mesopt., mesopterygium; metapt., metapterygium; neurocr., neurocranium; scapcor., scapulocoracoid; palatoq, palatoquadrate; propt., propterygium; R., right; vert., vertebrae; I-III, components of branchial arches one to three. All scale bars 5 mm.

### Manual preparation

The specimens were manually prepared at the NHMUK to extract one tooth from each. A combination of mechanical and chemical preparation was used to remove an isolated tooth from within the matrix for each specimen. Preparation was then carried out under a Leica M80 stereo microscope, with a ‘cube’ of matrix containing the tooth being excavated using a pin vise. The cube was then placed in a small container of 2% acetic acid using calcium orthophosphate as a buffer. The tooth was then bathed in water and cleaned with a fine bristle brush. The two isolated teeth, NHMUK PV P 17723A and NHMUK PV P 73821D, (Figure 3, Supplementary Figures 4, 5) were then imaged using scanning electron microscopy (SEM) on a JEOL JSM-IT500 scanning electron microscope at the NHMUK.

**Figure 3.**
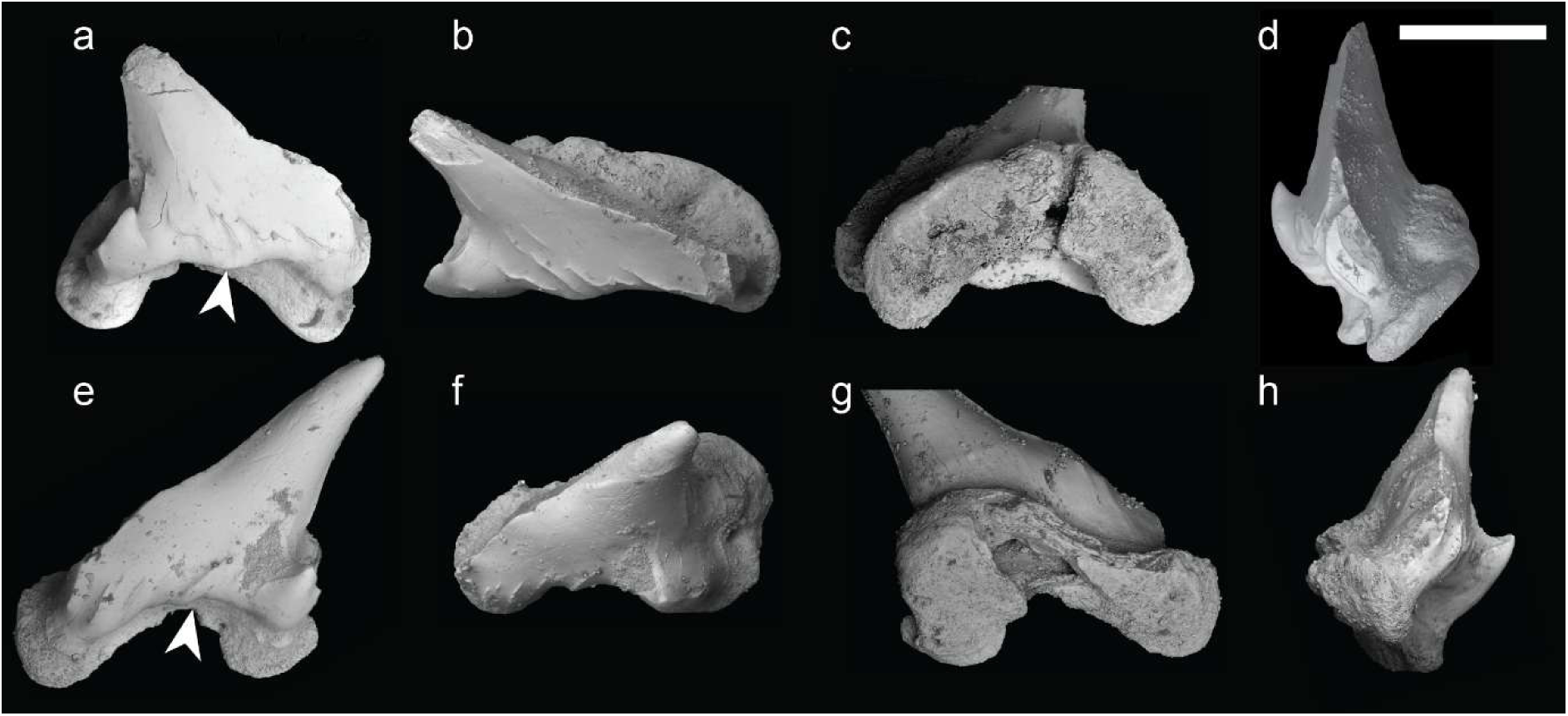
Scanning electron micrographs of the teeth of *Pararhincodon n. sp.* a-d isolated tooth of *P. n. sp.* NHMUK PV P 17723a in labial (a), occlusal (b), lingual (c), and anterior (d) views; e-h; isolated tooth of *P. n. sp.* NHMUK PV P 73821D in labial (e), occlusal (f), lingual, (g), and anterior (h) views. Arrow denotes medial protruberance. Scale bar = 0.5 μm.

### Phylogenetic analysis

We assembled a phylogenetic matrix with 19 taxa and 98 morphological characters for galeomorphs with a focus on carpet sharks, primarily from Goto (2001) but also using additional sources (see supplementary details for character list). We included representatives of major orectolobiform groups as well as other galeomorphs, and used two squaliforms as an outgroup. Details of taxa and phylogenetic characters are given in the Supplementary Materials.

We analysed the matrix using maximum parsimony and Bayesian approaches. Maximum parsimony analyses were carried out in Paup* 4.0 (Swofford, 2003); we performed a heuristic search using a TBR algorithm and 100,000 random addition sequence replicates, holding 5 trees at each step. Bayesian estimates of phylogeny were carried out in MrBayes 3.2.7a (Ronquist et al., 2012). The matrix was analysed using an Mkv model with gamma distributed variable rates and a uniform tree prior. We carried out two separate runs, each for 10,000,000 generations, and checked for convergence initially by checking the standard deviation of split frequency was below 0.01, and then by confirming adequate mixing in Tracer v1.7 (Rambaut et al., 2018) both visually and checking all ESS scores were >200.

We combined the trees from these runs using tools in R using the package [ape v5.8] (Paradis & Schliep, 2019) and calculated a majority rule consensus tree with a relative burn-in of 0.25.

## Results

### Systematic palaeontology

Class. **Chondrichthyes** Huxley, 1880

Subclass. **Elasmobranchii** Bonaparte, 1838

Superorder. **Galeomorphii** Compagno, 1973

Order. **Orectolobiformes** Applegate, 1972

Family. **Parascylliidae** Gill, 1862

Genus: ***Pararhincodon*** Herman in Cappetta, 1976

**Diagnosis** Extended from Cappetta 1980.

Parascylliid with elongate body. Neurocranium with rostral rod, no supraorbital crest, a mineralised cranial roof with an elongate epiphyseal foramen, a suborbital shelf, pointed postorbital processes, and widely-spaced endolymphatic ducts with no endolymphatic fossa. Long, narrow palatoquadrates with no posterior process. Meckel’s cartilage lacking posterior process. Vertebrae well-mineralised with primary calcification developed into four diagonal lamellae, notochordal space closed, and well-developed intermedialia. Anteriormost cervical vertebra broad with ventrolateral articulatory processes. Pectoral girdle approximately at level of twelfth vertebra, pro-, meso-, and metapterygia with strongly curved mesopterygium forming interspace with metapterygium.

**Remarks** This taxon shares with orectolobiforms an elongate rostral rod and the distinctive basipterygial interspace between the meso- and metapterygium. With parascylliids it shares the widely spaced endolymphatic ducts with no endolymphatic fossa, distinctively pointed postorbital processes, supraorbital crest absent, Meckel’s cartilage without posterior process, very small asymmetrical teeth. Unlike extant parascylliids the roof of the braincase and posterior orbital walls are mineralised, it has a separate propterygium in the pectoral fin, and the palatoquadrate lacks a posterior process. As in extant *Parascyllium* (*contra* Goto 2001) and in the asterospondylic vertebrae of extant carcharhiniforms the primary calcification of the vertebral centrae in *Pararhincodon* are developed into x-shaped diagonal lamellae.

Species: ***Pararhincodon*** n.sp.

**Derivation of name:** From torc, a metal collar associated with cultures from the European Iron Age.

**Holotype specimen:** NHMUK PV P 73821a

### Diagnosis of species

Very small, strongly asymmetrical teeth. Cusp triangular with sharp cutting edge, strongly bent lingually and distally. Lateral cusplet on distal edge of tooth, absent on mesial edge which is developed into slight shoulder. Base of cusp has a labial bulge and the medial section is developed into a low labial protuberance. Parallel folds present at the base of the cusp, some of which travel up the cusp’s face. Root flat and developed into mesial and distal lobes with deep, open nutrient groove.

**Remarks.** This new taxon differs from other *Pararhincodon* taxa in that the medial base of the labial crown is expanded into a low protuberance. The cusp is broader than in *P. crochardi*, and the folds resemble *P. ornatus*, although they are not developed into a horizontal crest (Guinot et al., 2013). Only lateral and posterior teeth are preserved, as asymmetry likely decreased towards the symphysis, a feature that is very pronounced in living parascylliids.

### Description

Both specimens present preserved cartilages three-dimensionally with minor crushing (Figure 2, Supplementary Figures 1, 2). Only calcified parts of the skeleton are preserved, i.e. the mineralised tesserae of the calcified cartilage and the areolar calcification of the vertebrae. All tesserae are single layered but there is variation in thickness, with tesserae on the mandibular arches being notably thicker than those of the neurocrania. The matrix has been bioturbated and numerous isolated dermal denticles (Supplementary Figure 6), are preserved in the matrix and in between the parts of the specimen as well as teeth, and pieces of invertebrates.

The neurocrania of both specimens is partially preserved, with the otic and occipital regions being most complete. The anterior edge of the neurocranial floor is preserved in NHMUK PV P 17223 curving forward to the fragmented rostral rod and base of the internasal septum and below the fragmentary posterolateral wall of the nasal capsule (Figures 2, 4a,e, Supplementary Figures 7, 8). The orbital wall is more extensively mineralised than in extant parascylliids, with tesselate cartilage surrounding what is likely a separate foramen for the oculomotor nerve (III) as well as foramina for the entry of the superficial ophthalmic + profundus nerve complex and the trigeminal and facial nerve (Figure 3, Supplementary Figures 7, 8). A supraorbital shelf is completely absent, with the orbital wall instead smoothly curving up onto the roof of the neurocranium, which is raised into a longitudinal hump like that in orectolobiforms such as *Chiloscyllium* (Staggl et al., 2022), and split by a long epiphyseal foramen. Unlike living parascylliids the neurocranial roof is calcified anteriorly beyond the level of the trigeminal/facial nerve foramen (Figure 4). A section of the left suborbital shelf is preserved in NHMUK PV P 17223 suggests that it was slightly more laterally extensive than in living parascylliids but the presence or absence of a foramen for the orbital artery or the position of the internal carotids entry into the basicranium are unknown (Figure 4a,d). As in *Parascyllium* the otic region features pronounced postorbital processes that taper to a point, and the endolymphatic openings are paired and broadly separated, lying posterolateral relative to the epiphyseal foramen, with no endolymphatic fossa or ascending process (Figure 4). Laterally to these the neurocranial roof is raised into low ridges over the anterior and posterior semicircular canals. Even allowing for some crushing, the postorbital region appears to be squatter than in *Parascyllium* with a more pronounced sphenopterotic ridge (Figure 4e,f). The hyomandibular facet is positioned immediately posteriorly to the orbit and is overhung by the triangular expansion of the postorbital process and is not set into the basicranial floor to the same extent as seen in *Parascyllium* (Figure 4). The postorbital process itself is large and triangular and the lateral surface of the otic capsule below it is slightly embayed. The posterior extent of the postorbital process is defined by the exterior semicircular canal, and below this slants downwards. The small internal foramen by which the glossopharyngeal nerve (IX) enters the cerebral cavity is visible in NHMUK PV P 73821a, and the nerve exits by its foramen at the posterior base of the postorbital process. The occipital region is broad with the foramen magnum flanked by foramina for the vagus (X) nerve (Supplementary Figure 7h). Ventral to it are broad paired occipital condyles and a median occipital hemicentrum.

**Figure 4.**
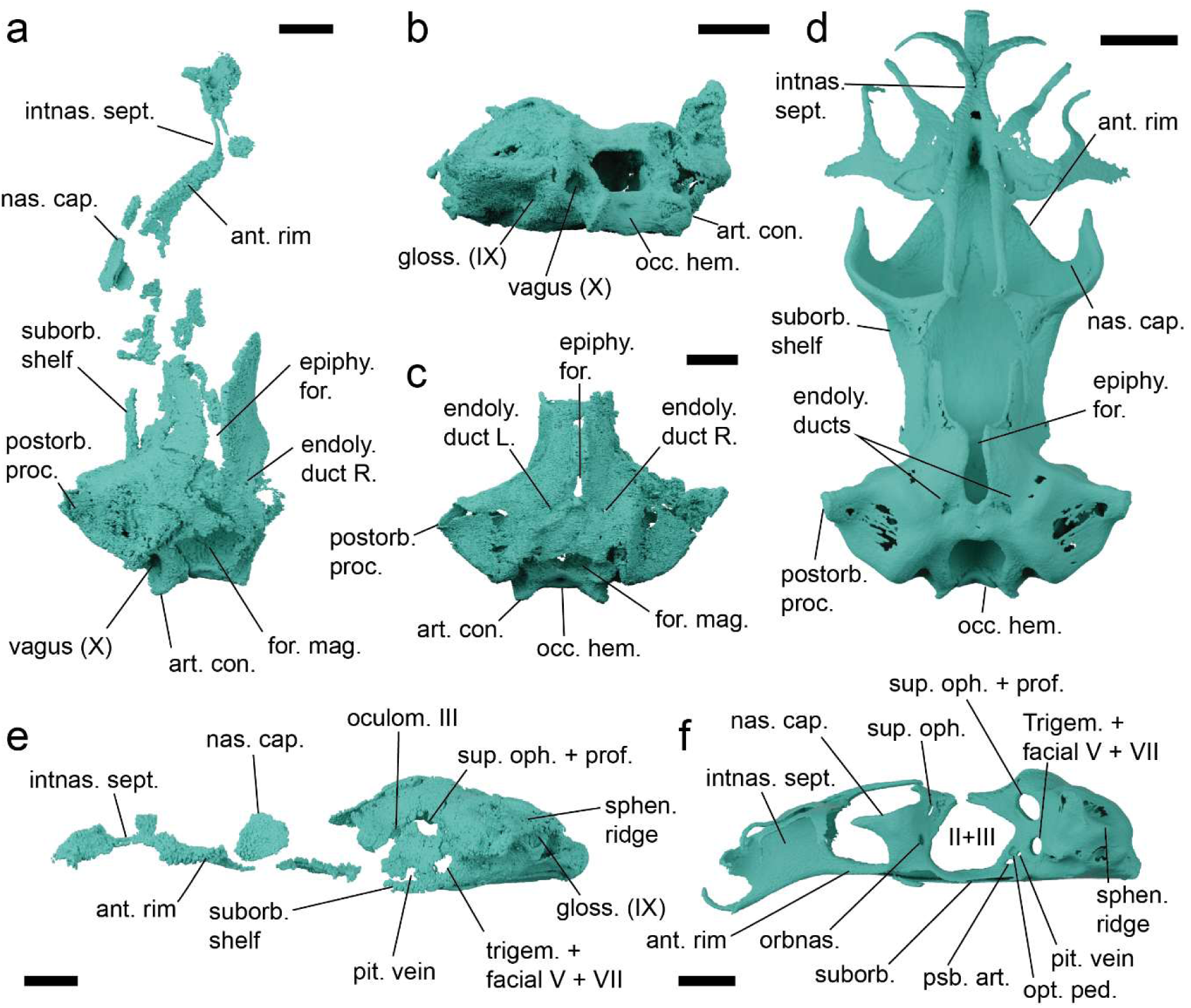
Details of the neurocranial anatomy of *Pararhincodon n. sp.* by comparison to the extant parascylliid *Parascyllium variolatum*. a, the neurocranium of *P. n. sp.* NHMUK PV P 17723 in dorsal view; b,c the neurocranium of *P. n. sp.* NHMUK PV P 73821a in postero- lateral (b) and dorsal (c) view; d, the neurocranium of *P. variolatum CSIRO* CA 3311 in dorsal view; e, the neurocranium of *P. n. sp.* NHMUK PV P 17723 in left lateral view; f, the neurocranium of *P. variolatum* CSIRO CA 3311in left lateral view. Abbreviations: ant. rim, anterior rim of basicranial floor; art. con., articular condyle; endoly., endolymphatic; epi. for., epiphyseal foramen; for. mag., foramen magnum; gloss. (IX), foramen for glossopharyngeal (IX) nerve; II+III, foramen for optic (II) and oculomotor (III) nerves; intnas. sept., internasal septum; nas. cap., nasal capsule; occ. hem., occipital hemicentrum; oculom. III, foramen for oculomotor (III) nerve; opt. ped., optic pedicel; pit., pituitary; postorb. proc., postorbital process; psb. art., foramen for pseudobranchial artery; suborb., suborbital; sphen. ridge, sphenotic ridge; suborbital; sup. oph + prof., foramen for superficial ophthalmic and profundus nerve; trigem. + facial V + VII; foramen for trigeminal and facial nerves; vagus (X), foramen for vagus (X) nerve. All scale bars 5 mm.

The jaws of both specimens are preserved and resemble those of *Parascyllium*. Palatoquadrates are straight with a low profile (Figure 5, Supplementary Figure 9) with a dental sulcus, low ethmo-palatine process and a broad ridge along the posterodorsal edge creating the mandibular adductor fossa. The posterior end of the palatoquadrate is blunt and developed into an angle, but lacks the posterior process present in extant parascylliids (Goto, 2001) (Supplementary Figure 9). With a concavity in the lingual surface forming the articulation with the Meckel’s cartilage. The Meckel’s cartilage is straight (Figure 5, Supplementary Figure 10) with a pronounced mandibular knob and a broad lateral flange carrying the articular cotylus. The posterior margin of both Meckel’s cartilages is preserved in NHMUK PV P 73821a and shows it did not bear a posterior process (Figure 5c,e). No labial cartilages are preserved in either specimen.

**Figure 5.**
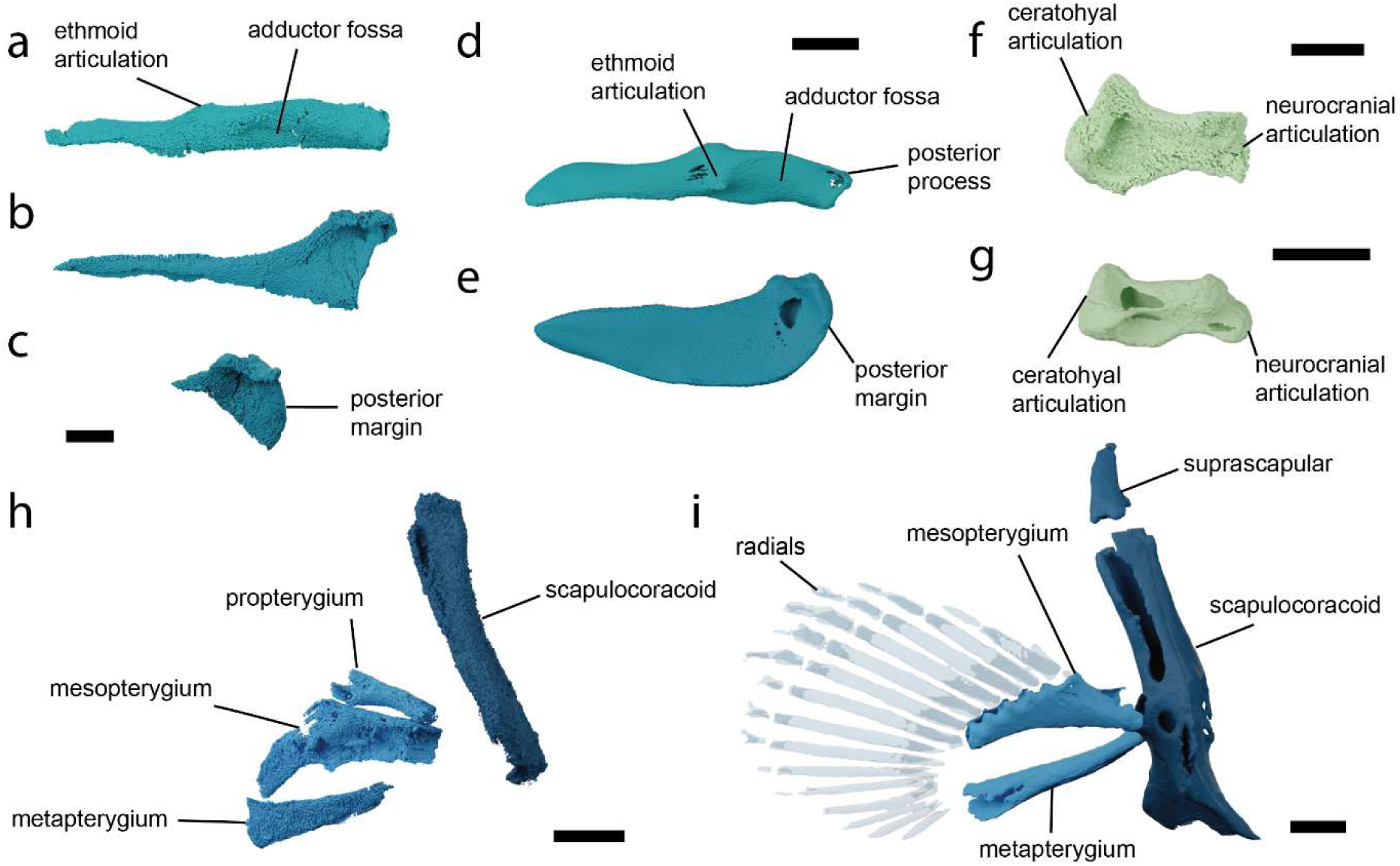
Details of the visceral and pectoral skeleton of *Pararhincodon n. sp.* by comparison to the extant parascylliid *Parascyllium variolatum*. a, the left palatoquadrate of *P*. *n. sp.* NHMUK PV P 17723 in labial view; b, the Meckel’s cartilage of *P. n. sp.* NHMUK PV P 17723 in labial view; c the articular region of Meckel’s cartilage of *P. n. sp.* NHMUK PV P 73821a in labial view; d, the palatoquadrate of *P. variolatum CSIRO* CA 3311 in labial view; e, the Meckel’s cartilage of *P. variolatum* CSIRO CA 3311 in labial view; f, the left hyomandibula of *P. n. sp.* NHMUK PV P 17723 in dorsal view, g, the left hyomandibula of *P. variolatum CSIRO* CA 3311 in dorsal view; h, right pectoral fin skeleton of *P. n. sp.* NHMUK PV P 73821a arranged into approximately life position in right lateral view; i, right pectoral fin skeleton of *P. variolatum* CSIRO CA 3311 in right lateral view. All scale bars 5 mm.

Although no teeth are preserved in articulation with the mandibular cartilages in either specimen, we interpret teeth in both specimens as originating from the articulated specimens due to their consistent morphology and size and their location close to the mandibular cartilages (Figure 3, Supplementary Figures 4,5). In NHMUK PV P 73821a we identify nine teeth in the matrix and in NHMUK PV P 17223 four teeth (Supplementary Figure 5). Teeth are very small, being approximately 1 mm in height (Figure 3, Supplementary Figure 4). All preserved teeth are strongly asymmetrical and represent lateral or posterior teeth (Cappetta, 1987, 2012), which is consistent with both specimens missing the front part of the jaws. A detailed description of tooth morphology is given in the diagnosis.

The full length of the pharynx is preserved in NHMUK PV P 73821a, and parts of the hyoid skeleton are preserved in both specimens. The hyomandibula is short and robust, with the distal surface expanded for its articulation with the palatoquadrate/ceratohyal to a greater extent than in extant *Parascyllium* (Figures 2, 5, Supplementary Figure 11). The ceratohyal is arched and broad, with a low groove on the labial surface (Figure. 2, Supplementary Figure 12). The basihyal is missing in both specimens. The branchial skeleton is preserved in both specimens but more extensively in NHMUK PV P 73821a (Figure 2, Supplementary Figure 13). In NHMUK PV P 17223 the anteriormost left branchial arch is preserved, including most of ceratobranchial I, epibranchial I and the proximal tip of pharyngobranchial I. In NHMUK PV P 73821a the branchial skeleton is heavily disrupted but on the left side three epibranchials (likely I-III) are preserved, while on the right side there are several branchial cartilages including identifiable cerato-, epi-, and pharyngobranchials. Also preserved is the tip of the anterior process of the basibranchial copula. The branchial cartilage proportions and morphology match those of *Parascyllium* (Figure 2, Supplementary Figure 13).

The pectoral skeleton is preserved only in NHMUK PV P 73821a (Figures 2d,, Supplementary Figure 14). Flattening of the shoulder girdle during preservation has resulted in the dorsal processes of the scapulocoracoids being pushed posteriorly relative to the fin skeleton and shattered pieces of cartilage are likely corresponding to parts of the articular regions and the coracoid bar. The basipterygia of both left and right pectoral fins are preserved. Meso- and metapterygia are identifiable by comparison to those of *Parascyllium*, although they have slightly different proportions. Associated with these are additional elements: based on their morphology we are able to eliminate the possibility that they are posterior branchial elements, and based on their location they are unlikely to be superscapulae. We conclude that these are propterygia, which are absent in extant parascylliids (Goto, 2001). No fin radials are preserved.

Sections of the vertebral column are preserved in both specimens: NHMUK PV P 17723 preserves the anteriormost three vertebral centra while NHMUK PV P 73821a preserves a largely articulated section of the vertebral column extending to the level of the pectoral girdle, comprising 18 vertebral centra (Figures 2, 6, Supplementary Figure 15). Interdorsal and basidorsal elements are also present but mostly scattered through the matrix (Supplementary Figure 2). Vertebrae comprise cone-shaped zones of primary calcification, with the notochordal space either absent or very small. Cross-sections of the vertebrae of *Pararhincodon* from throughout the preserved column show that the primary calcification extends via four short mineralised lamellae, arranged into an x-shape, into the intermedialia (Figure 6a,b). Although this fits within the range of conditions described as ‘asterospondylic’ (Cappetta 1987), the numerous secondary calcifications that extend radially into the intermedialia in most orectolobiforms, either to the level of the *membrana elastica externa* (hemiscylliids, *Stegostoma*, *Ginglymostoma*, Figure 6c) or not reaching this level (orectoloboids and *Brachaelurus*), are absent (Figure 6d-f). The outer surfaces of the vertebrae are well mineralised. The anteriormost vertebral centrum has expanded processes for articulation with the occipital condyles. *Contra* Goto (2001), who described the vertebrae of living parascylliids as lacking any kind of extension from the primary calcification our CT-scans of *Parascyllium* show that these are present throughout the vertebral column and strongly resemble *Pararhincodon* (Figure 6c). Similar x-shaped laminae extending from the primary calcification are also present in Carcharhiniformes (Cappetta, 1987; Ridewood, 1921; White, 1937), although parascylliids lack the Maltese-cross shaped calcified areas in taxa like *Prionace* (Figure 6e) instead more closely resembling the vertebrae of *Proscyllium* (Figure 6f).

**Figure 6.**
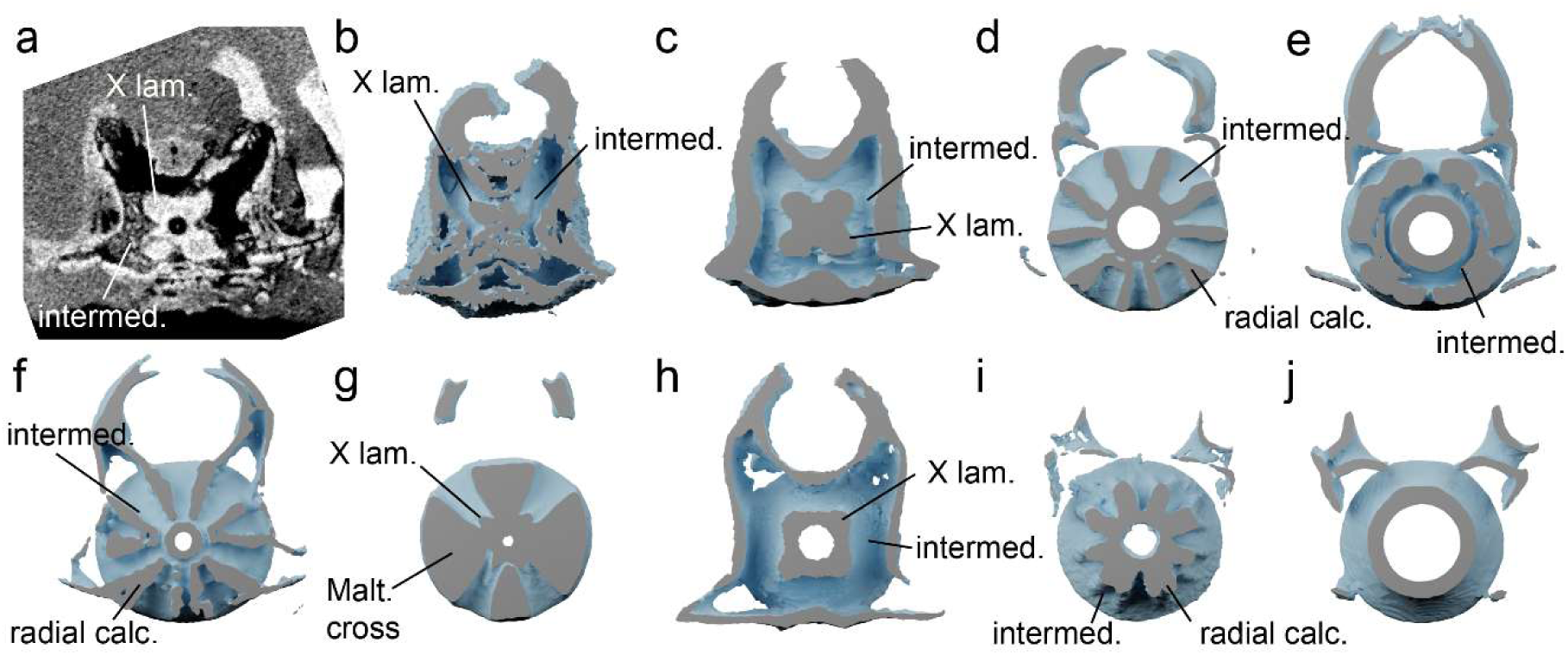
Bisected vertebrae of *Pararhincodon n. sp.* by comparison to extant sharks, showing mineralised regions based segmented from CT data. All vertebrae taken from the level of approximately the heart. a,b, *P. n. sp.* NHMUK PV P 73821a. as a CT section (a) and bisected 3D volume (b); c, *Parascyllium variolatum CSIRO* CA 3311; d, orectolobiform *Chiloscyllium plagiosum*; e, orectolobiform *Sutorectus tentaculatus*; *f*, orectolobiform *Brachaelurus waddi;* g, carcharhiniform *Prionace glauca*; h, carcharhiniform *Proscyllium magnificum*; i, heterodontid *Heterodontus francisci*; j, squaliform *Squalus cubensis*. Abbreviations: intermed., intermedialia; Malt. cross, Maltese-cross-shaped secondary calcification; rad. calc., radial secondary calcification; X lam, X-shaped laminae. Not to scale. See Supplementary Table 1 for specimen details. Not shown to scale.

### Phylogenetic analysis

The parsimony analysis recovered eight most parsimonious trees, with a length of 192. Both Bayesian and parsimony approaches recovered tree topologies broadly consistent with estimates of selachian phylogeny based on molecular data (Marion et al., 2024; Naylor et al., 2012). Galeomorphs were found to be monophyletic, with *Heterodontus* as the sister group to all other galeomorphs. Within galeomorphs, orectolobiforms are found to be the sister group to carcharhiniforms plus lamniforms. Our dataset recovered carcharhiniforms as being paraphyletic relative to lamniforms, likely as a result of poor taxon sampling.

Both analytical approaches recover orectolobiforms as a clade, with extant parascylliids, and hemiscylliids, *Stegostoma*, *Rhincodon*, and *Ginglymostoma* as sub-clades, reflecting molecular-based understanding of the group’s intrarelationships. *Brachaelurus* and wobbegongs (*Sutorectus*, and *Orectolobus*) are recovered in a polytomy with these clades: although unresolved this is consistent with molecular trees which suggest they form a clade that is the sister group to all other orectolobiforms except parascylliids (Marion et al., 2024; Naylor et al., 2012). Both approaches found *Pararhincodon n. sp.* to be a stem-group parascylliid within orectolobiforms, i.e. the sister group to a clade formed by *Parascyllium* and *Cirrhoscyllium*, with a high degree of confidence (Figure 7).

**Figure 7.**
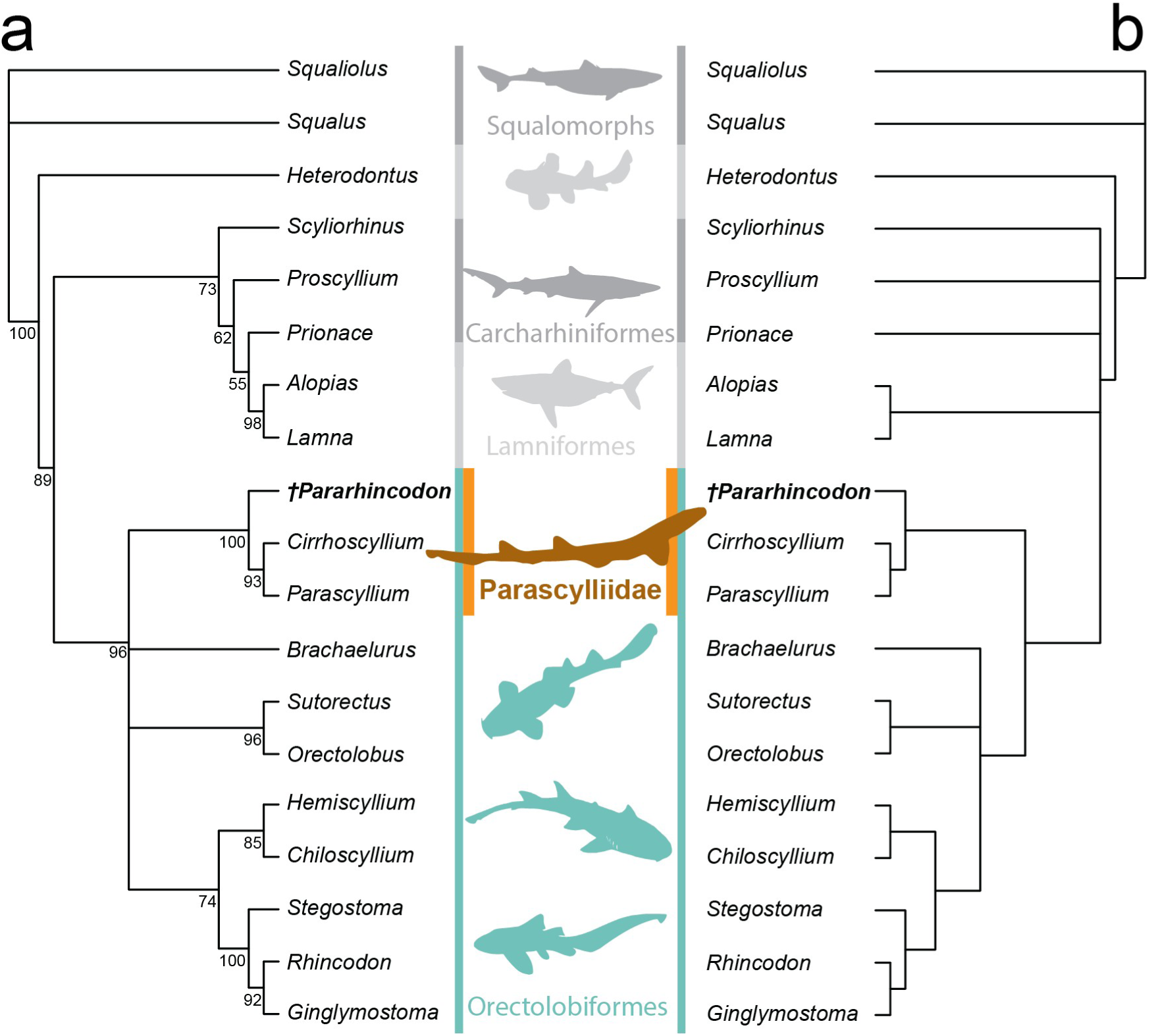
Results of phylogenetic analysis of morphological phylogenetic dataset. a, 50% Majority rule consensus tree from Bayesian analysis with node values representing posterior probability of that clade; b, strict consensus of eight most parsimonious trees recovered by parsimony analysis. Silhouettes from Phylopic: see Supplementary Table 2 for attributions.

## Discussion

### *Pararhinchodon* and the Parascylliidae

Based on our analysis we are able to confidently identify *Pararhincodon n. sp.* as a stem-group parascylliid (Figure 7). Previously, articulated *Pararhincodon* were known only from Lebanon (Pfeil, 2021), and to date only one species, *Pararhincodon lehmani* from the Cenomanian of Haqel, was formally described based on articulated remains (Cappetta, 1980). This description was based on a single poorly preserved specimen, MNHN HAK 556, which was attributed to Orectolobiformes on the basis of the presence of a rostral rod on the neurocranium, and then to the Parascylliidae on the basis of teeth lacking a labial apron (Cappetta, 1980). As a result, *Pararhincodon*’s status as a parascylliid has been treated cautiously (Pfeil, 2021) and the taxon has been omitted from attempts to reconstruct orectolobiform evolutionary history (Boyd & Seitz, 2021). In addition to *Pararhincodon lehmani* and *P*. *n. sp.*, three species based on isolated teeth are known ranging from the Cenomanian to the Lutetian - *P. crochardi*, *P. germaini*, and *P. ypresiensis* (Adnet, 2006; Cappetta, 1976, 2012; Herman, 1977) and the oldest teeth assigned to the genus are Albian (Underwood & Mitchell, 1999). Our robust identification of *Pararhincodon* as a parascylliid using skeletal morphology allows this tooth record to be associated with the parascylliid stem-group with confidence.

Extant parascyllliids comprise two genera, *Parascyllium* and *Cirrhoscyllium*, which molecular and morphological data indicate are the sister-group to all other orectolobiforms (Boyd & Seitz, 2021; Goto, 2001; Marion et al., 2024). These taxa are distinguished from other orectolobiforms by numerous distinctive skeletal anatomies of the neurocranium, visceral skeleton and pectoral girdle (Goto, 2001), and have a markedly anguilliform body shape which within orectolobiforms is comparable only to some hemiscylliids (Ebert et al., 2013).

Our new data show that much of this skeletal anatomy, for example the distinctive morphology of the otic region of the neurocranium and the internal mineralization of the vertebrae (Goto, 2001), were already in place well before the divergence of the crown-group. However, other aspects, such as the loss of the propterygium in the pectoral fin and demineralisation of the neurocranium, were more recently acquired and may be features of the parascylliid crown-group. Comparison to the body profile of undescribed *Pararhincodon* specimens from Haqel and Hjoula (Pfeil, 2021, pl. 20), suggests that the body of *Pararhincodon* was less anguilliform with more anteriorly placed dorsal fins than extant parascylliids. Contemporaneous Lebanese parascylliids show that the more pronouncedly anguilliform body shapes in parascylliids had already evolved by this point and these may represent closer relatives of the parascylliid crown-group (Pfeil, 2021). These new data provide an insight into the mosaic evolution of modern parascyllids’ distinctive anatomies.

The divergence of *Parascyllium* and *Cirrhoscyllium* is comparatively recent, placed by molecular estimates in the Late Cretaceous or Palaeogene (Boyd & Seitz, 2021; Marion et al., 2024), with the oldest tooth assigned to a living genus (*Parascyllium*) from the Lutetian (Eocene) of France (Adnet, 2006; Cappetta, 2012). Their divergence from other orectolobiforms is much older than this date or the range of *Pararhincodon,* with molecular estimates placing it in the Triassic (Boyd & Seitz, 2021; Marion et al., 2024). Spanning this gap by confidently assigning Jurassic taxa to the parascylliid stem-group is challenging due to the small size and fairly unremarkable anatomy of parascylliid teeth, illustrated by the initial assignment of *Pararhincodon* to the Scyliorhinidae (Cappetta, 1976). In most *Pararhincodon* species a labial apron is absent, however the labial protuberance on the teeth of *Pararhincodon n. sp.* bears comparison with the distinctive apron of orectolobiform teeth and suggests that this could be a plesiomorphic feature of their dentitions rather than a feature uniting other Orectolobiformes to the exclusion of parascylliids. If so, candidate stem-group parascylliids are not limited to taxa with tooth aprons, for example *Phorcynis,* which has a larger labial protuberance than *Pararhincodon n. sp.* but a similarly asymmetrical shape of the crown (Cappetta, 2012). Other Mesozoic tooth taxa have crown morphologies that depart further from that of *Pararhincodon* but possess labial protuberances and slender cusps, for example *Akaimia* (Rees, 2010; Srdic et al., 2016).

### Orectolobiformes and galeomorphs

Because the divergence of parascylliids from other orectolobiforms is equivalent to the divergence of the orectolobiform crown-group the timescale of their evolution has important implications for the evolution of the galeomorph crown-group. Various holomorphic Upper Jurassic fossils have been identified as Orectolobiformes including *Phorcynis*, *Palaeorectolobus* and possibly *Palaeoscyllium* (Kriwet, 2005; Villalobos-Segura et al., 2023) while the oldest isolated teeth assigned to Orectolobiformes are Toarcian examples of *Annea* and *Folipistrix* (Cappetta, 2012; Delsate, 2003; Delsate & Thies, 1983; Kriwet, 2003). However, a lack of phylogenetically informative characters means that these Jurassic taxa, whether teeth or holomorphic specimens, cannot at present be assigned to specific internal branches of the orectolobiform tree. Recent molecular estimates place major crown-group orectolobiform divergence events, i.e. the split between the clade including Brachaeluridae and Orectolobidae and the clade including Hemiscylliidae, Ginglymostomatidae, and Rhincodontidae (Figure 7), as well as early divergences within those clades, in the Jurassic/Lower Cretaceous. Jurassic Orectolobiformes could feasibly represent stem-group members of those lineages, but could also feasibly represent stem-group Orectolobiformes. Although it is considerably younger than these taxa, strictly speaking the Cenomanian *Pararhincodon n. sp.* is a candidate for a robust minimum constraint for the orectolobiform crown-group because it can be confidently assigned to a specific branch of the crown-group (Parham et al., 2012). Less conservatively the Campanian putative ginglymostomatid *Cantioscyllium decipiens* could also provide a hard minimum constraint for the orectolobiform crown-group, and *Cantioscyllium* teeth have been used to constrain this node in recent time-calibrated phylogenetic analysis (Marion et al., 2024).

The similarities in vertebral anatomy that we identify between parascylliids and carcharhiniforms fit into an emerging understanding of elasmobranch macroevolution. Galeomorphs’ long evolutionary history probably extends back to the Permian, and extant galeomorph groups are thought to have evolved in coastal habitats from relatively small, benthic forms (Marion et al., 2024; Sorenson et al., 2014; Sternes et al., 2024). The presence of four diagonal lamellae themselves probably form the basis for more complex asterospondylic vertebral conditions across elasmobranchs during development (Ridewood, 1921).However, the adult parascylliid-like configuration differs from all other orectolobiforms including asterospondylic forms (Goto, 2001) and instead is strikingly similar to that of several catshark-like carcharhiniform taxa including *Proscyllium* and *Schroederichthys* (Ridewood, 1921; White, 1937). No Lamniformes display these x-shaped laminae, instead having vertebrae with numerous radial lamellae (Knaub et al., 2024; Ridewood, 1921). Thus, this x-shaped configuration could plausibly represent a plesiomorphic anatomy for galeomorphs excluding Heterodontiformes, retained in certain crown-group members but lost in disparate members of the group. Its retention in adult parascylliids and catshark-like carcharhiniforms may reflect the lower mechanical demands of small body size and benthic lifestyles, with larger galeomorphs supporting larger body and life in the water column by increasing the level of secondary calcification within the intermedialia. This might also explain this state’s absence in lamniforms; while some early lamniform teeth are found in nearshore environments all living lamniforms are large and free-swimming (Kriwet et al., 2008; Rees, 2005; Sternes et al., 2024).

Finally, these new data show the potential of shark skeletons preserved in the Chalk as a window into the evolution of modern elasmobranch biodiversity. Modern orectolobiform biodiversity is centred in the Indo-Pacific biodiversity hotspot, and most species-level orectolobiform biodiversity originated in this region during the Cenozoic (Boyd & Seitz, 2021; Dudgeon et al., 2020). The two extant parascylliid genera are highly geographically restricted within this region, with *Parascyllium* endemic to Australia and *Cirrhoscyllium* to Vietnam, Taiwan, and Japan (Goto & Last, 2001; Goto & Nakaya, 1996). However, the fossil record indicates that this biodiversity centre shifted from the shallow seas of the Atlantic in the Upper Cretaceous to its present location via the Tethys Sea through the Cenozoic, in the wider context of a global shift in marine biodiversity (Boyd & Seitz, 2021; Renema et al., 2008). As our description of *Pararhincodon n. sp.* shows, the Chalk has the potential to provide a uniquely detailed glimpse into elasmobranch faunas at the beginning of this process, and to complement and be complemented in turn by the detailed picture provided by the tooth record (Guinot et al., 2013). Further descriptions of crown-group elasmobranchs from the Chalk should allow the incorporation of several taxa confidently into elasmobranch phylogeny, and estimates of their evolutionary timing.

## Conclusions

Three-dimensional fossil remains of crown-group elasmobranch skeletons are much rarer than their teeth but are far more information-rich. As it does for actinopterygians (Beckett et al., 2017; Friedman et al., 2015) the Cretaceous Chalk provides a uniquely detailed insight into these skeletal anatomies, allowing the reconciliation of the records of isolated teeth and skeletal morphologies which are poorly understood even in those elasmobranchs known from exquisitely preserved holomorphic fossils (Pfeil, 2021; Villalobos-Segura et al., 2023). Here, the skeletal anatomy of the Cenomanian *Pararhincodon n. sp.* allows the confident identification of *Pararhincodon* as a stem-group parascylliid and provides a rare insight into the morphological transformations that led to modern orectolobiform biodiversity. The Chalk offers a unique opportunity to harvest skeletal information from extinct elasmobranchs which should ultimately feed morphological information into elasmobranch phylogeny, estimates of the timing of their evolution, and efforts to understand phenomena like their slow evolutionary rate (Sendell-Price et al., 2023).

## Acknowledgements

Thanks to the NHMUK Imaging and Analysis Centre for scanning, particularly Dr Alex Ball (NHMUK, London) for assistance with SEM and Vincent Fernandez and Agnese Lanzetti for assistance with micro-CT scanning fossils at NHMUK. The authors acknowledge the facilities and scientific and technical assistance of the National Imaging Facility, a National Collaborative Research Infrastructure Strategy (NCRIS) capability, and Dr Ekaterina Strounina at the Centre for Advanced Imaging, University of Queensland for scanning the *Parascyllium variolatum* specimen. Thanks also to Guillaume Guinot, Sebastian Stumpf, and Eduardo Villalobos-Segura for helpful discussion. Many thanks also to Dr Brian Gasson who donated NHMUK PV P 73821 to the NHMUK collection. Finally, we would like to thank Jürgen Kriwet and one additional anonymous reviewer who provided helpful comments on the manuscript.

## Supplementary Data

All data supporting this study will be made freely available on publication of the final paper.

## Supplementary Figure legends

**Supplementary Figure 1.**
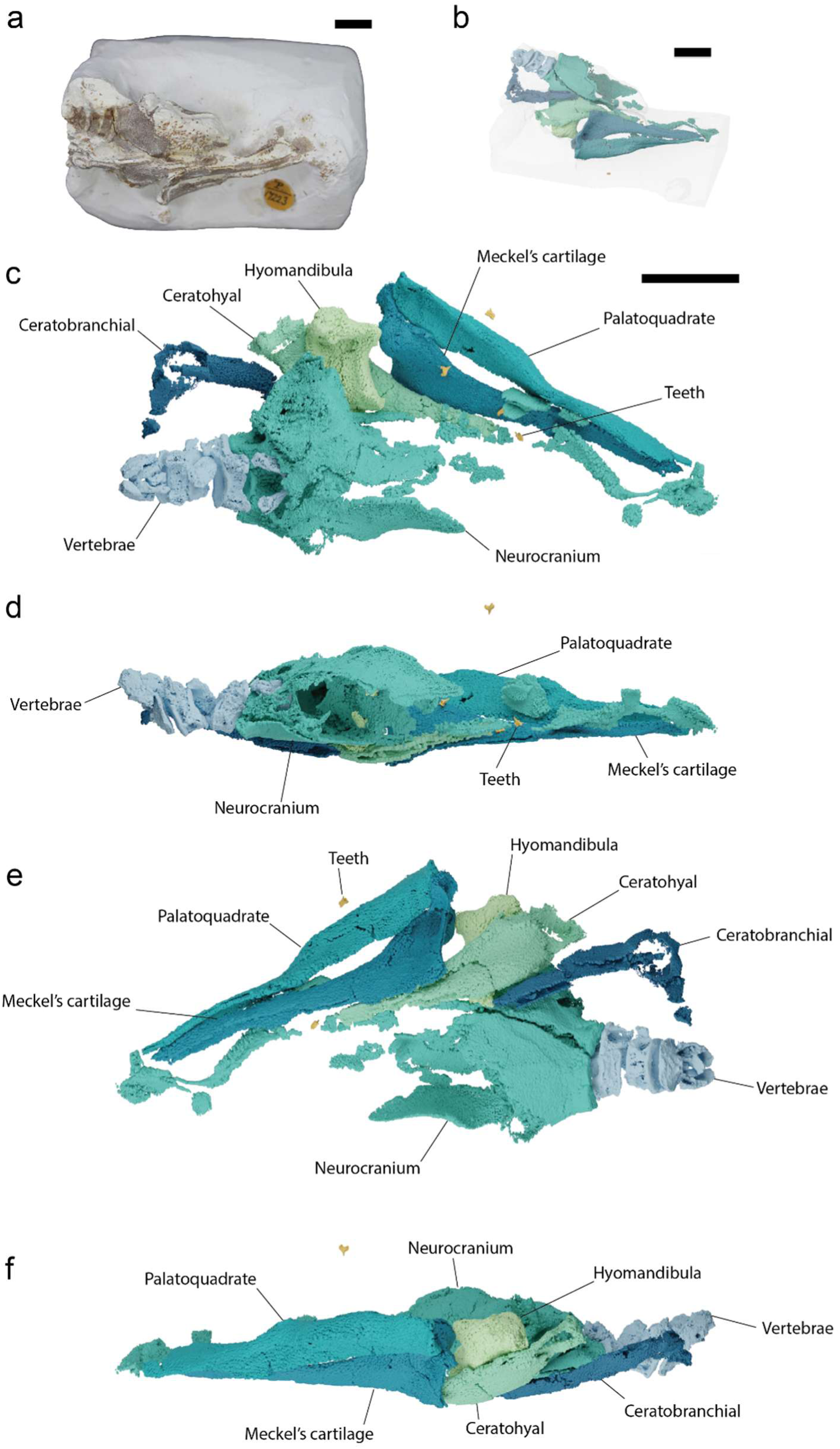
*Pararhincodon n. sp.* NHMUK PV P 17223. a, photograph of specimen; b, tomographic model showing positions of skeleton in matrix; c-f, 3D models of skeleton in dorsal (c), right lateral (d), ventral (e), and left lateral (f) view. All scale bars 10 mm.

**Supplementary Figure 2.**
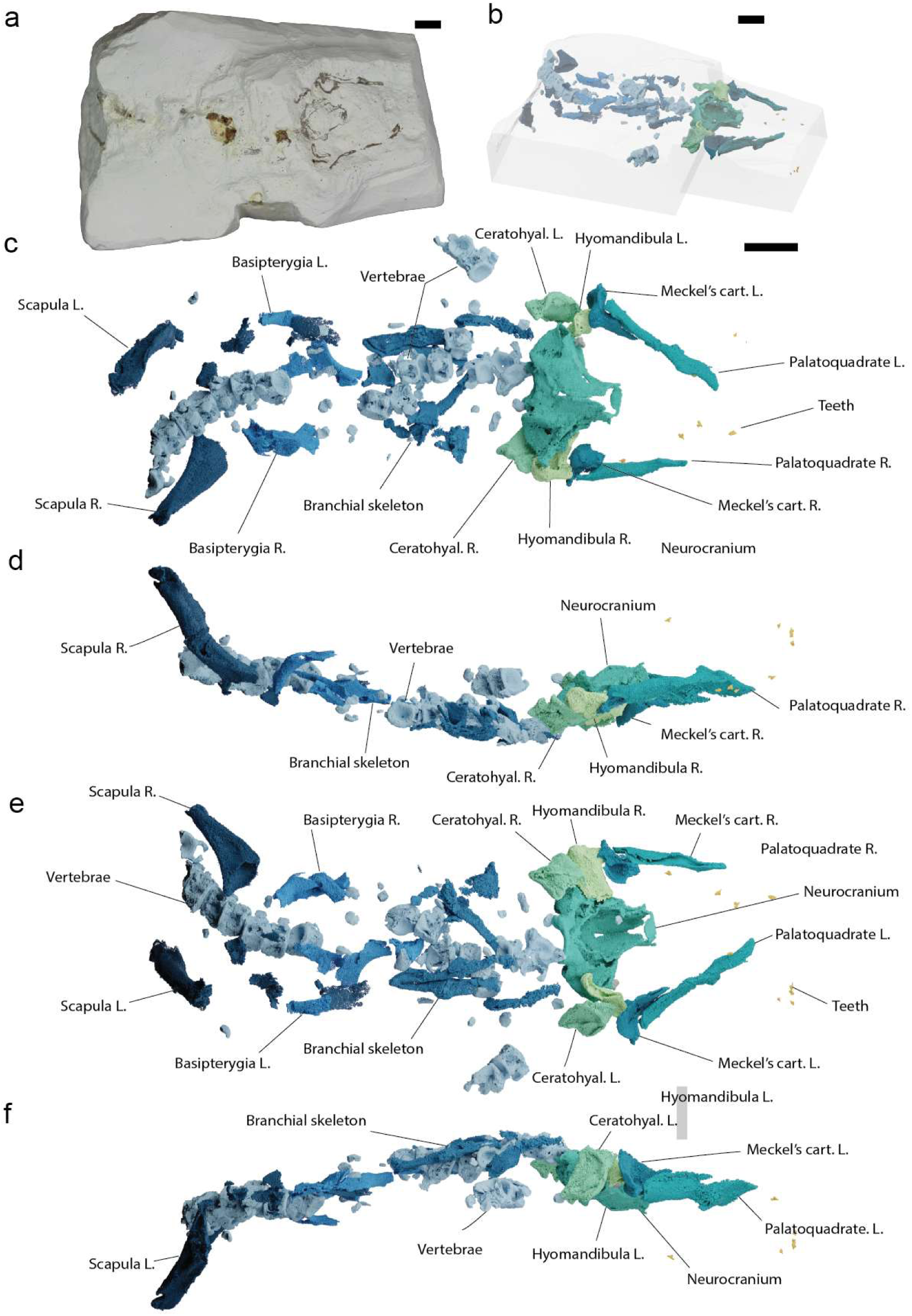
*Pararhincodon n. sp.* NHMUK PV P 73821a. a, photograph of specimen; b, tomographic model showing positions of skeleton in matrix; c-f, 3D models of skeleton in dorsal (c), right lateral (d), ventral (e), and left lateral (f) view. Abbreviations: L, left; R, right. All scale bars 10 mm.

**Supplementary Figure 3.**
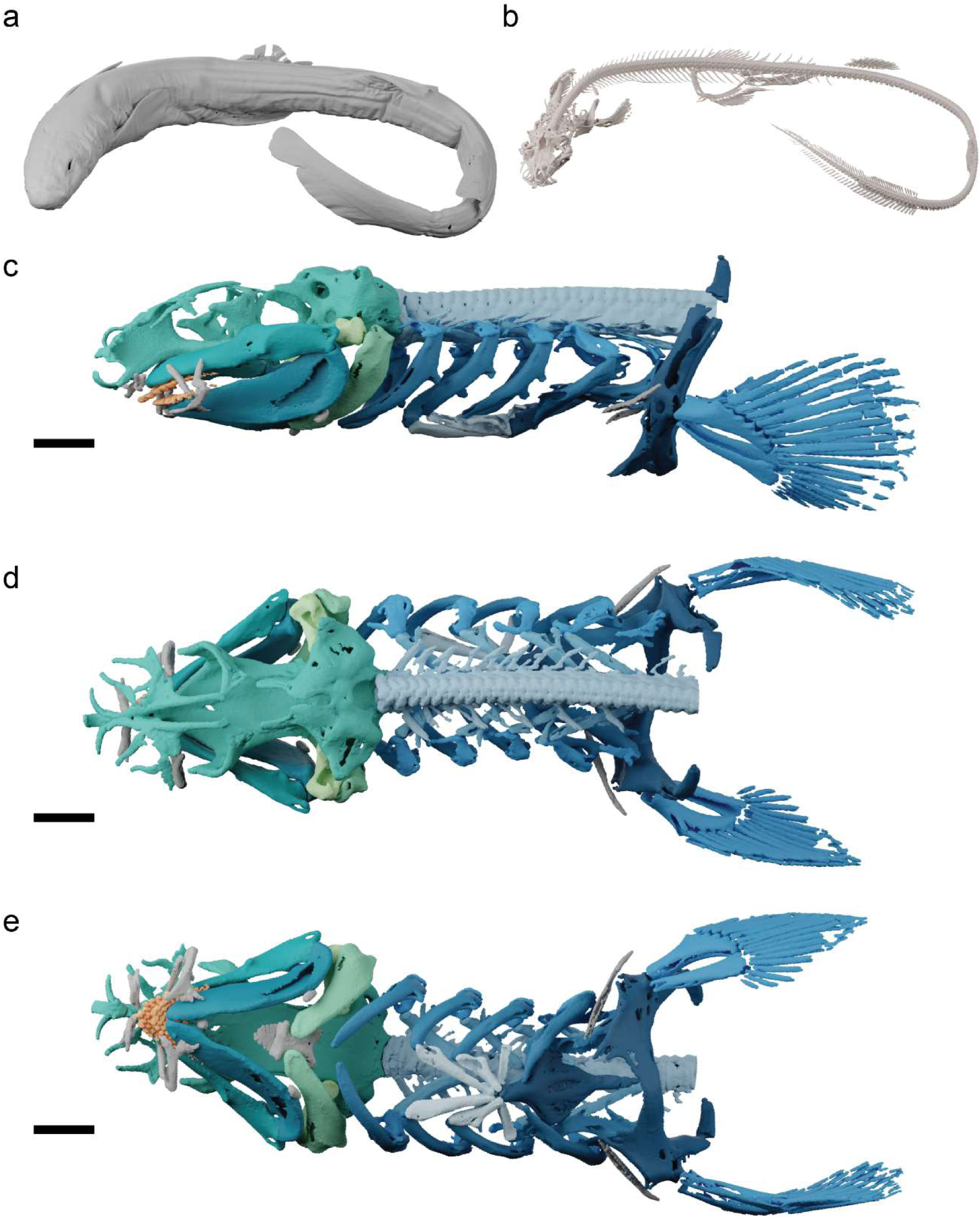
CT renders of *Parascyllium variolatum* CSIRO CA 3311. a, exterior of specimen; b, endoskeleton; c-d head and pectoral skeleton in left lateral (c), dorsal (d), and ventral (e) views. Scale bar 10 mm.

**Supplementary Figure 4.**
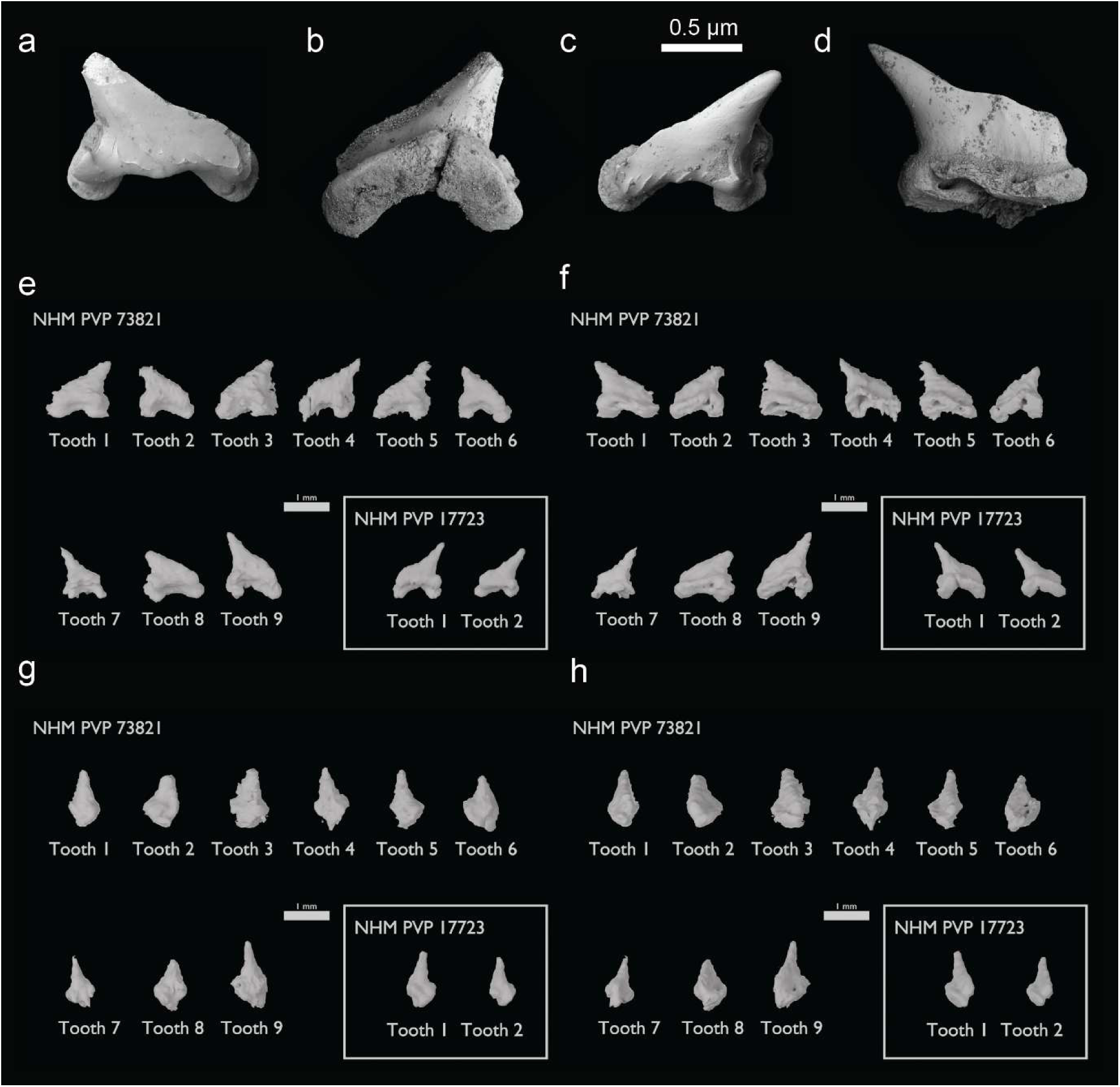
Additional images of the teeth of *Pararhincodon n. sp.* a,b isolated tooth *P. n. sp.* NHMUK PV P 17723a in occlusal-labial (a), and occlusal-lingual (b) views; c,d, isolated tooth *P. n. sp.* NHMUK PV P 73821d in occlusal-labial (c), and occlusal- lingual (d) views; e-f renders of teeth of both specimens from CT data in labial (e), lingual (f), posterior (g), and anterior (h) views.

**Supplementary Figure 5.**
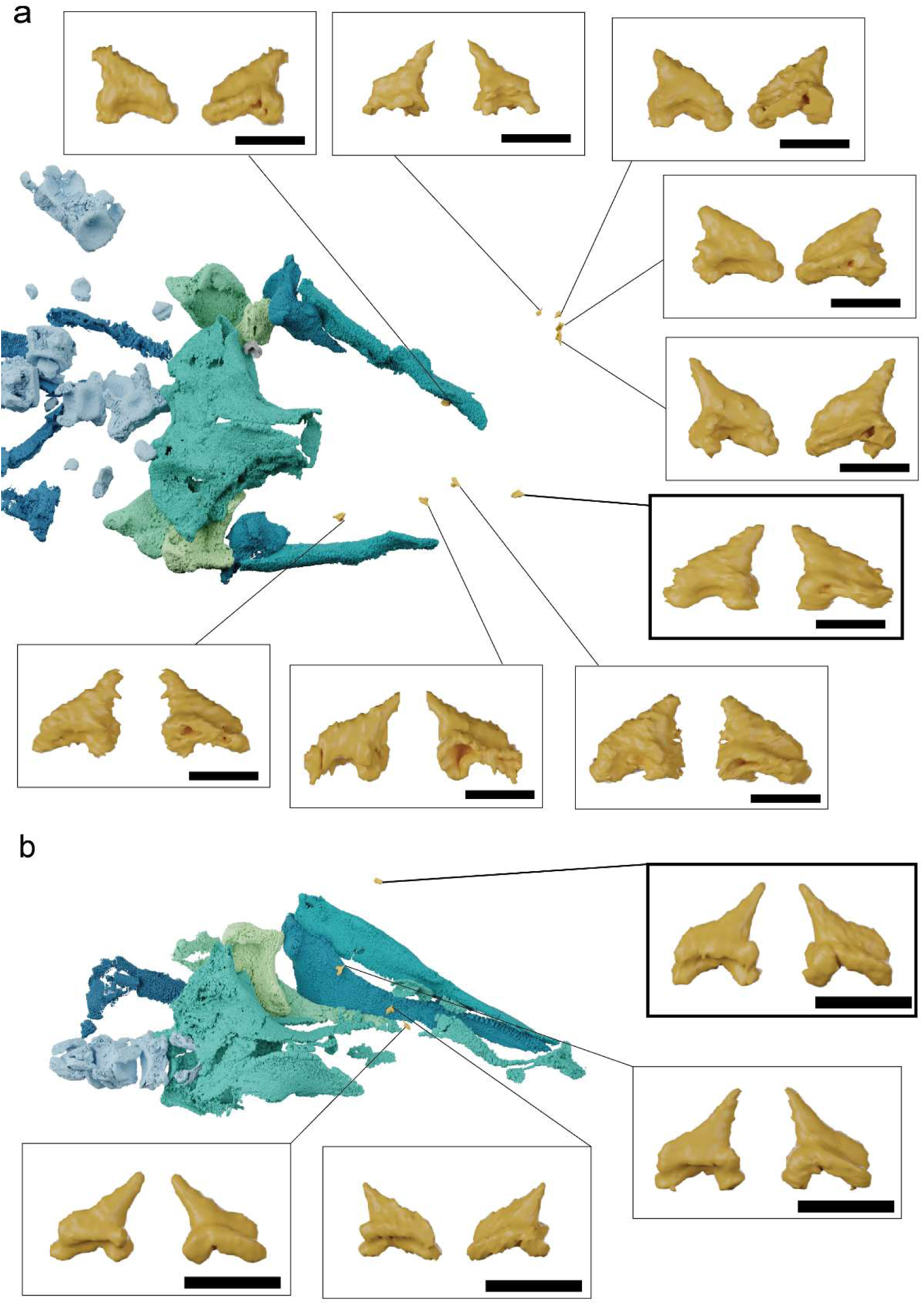
CT renders of teeth from both specimens of *Pararhincodon n. sp.* with enlarged insets showing the gross morphology of each tooth in lateral (left) and lingual (right) view: a, NHMUK PV P 73821a; b, NHMUK PV P 17223. Bold boxes indicate the teeth that were extracted for SEM, i.e .in panel a NHMUK PV P 73821d and panel b NHMUK PV P 17223a. All scale bars 1 mm.

**Supplementary Figure 6.**
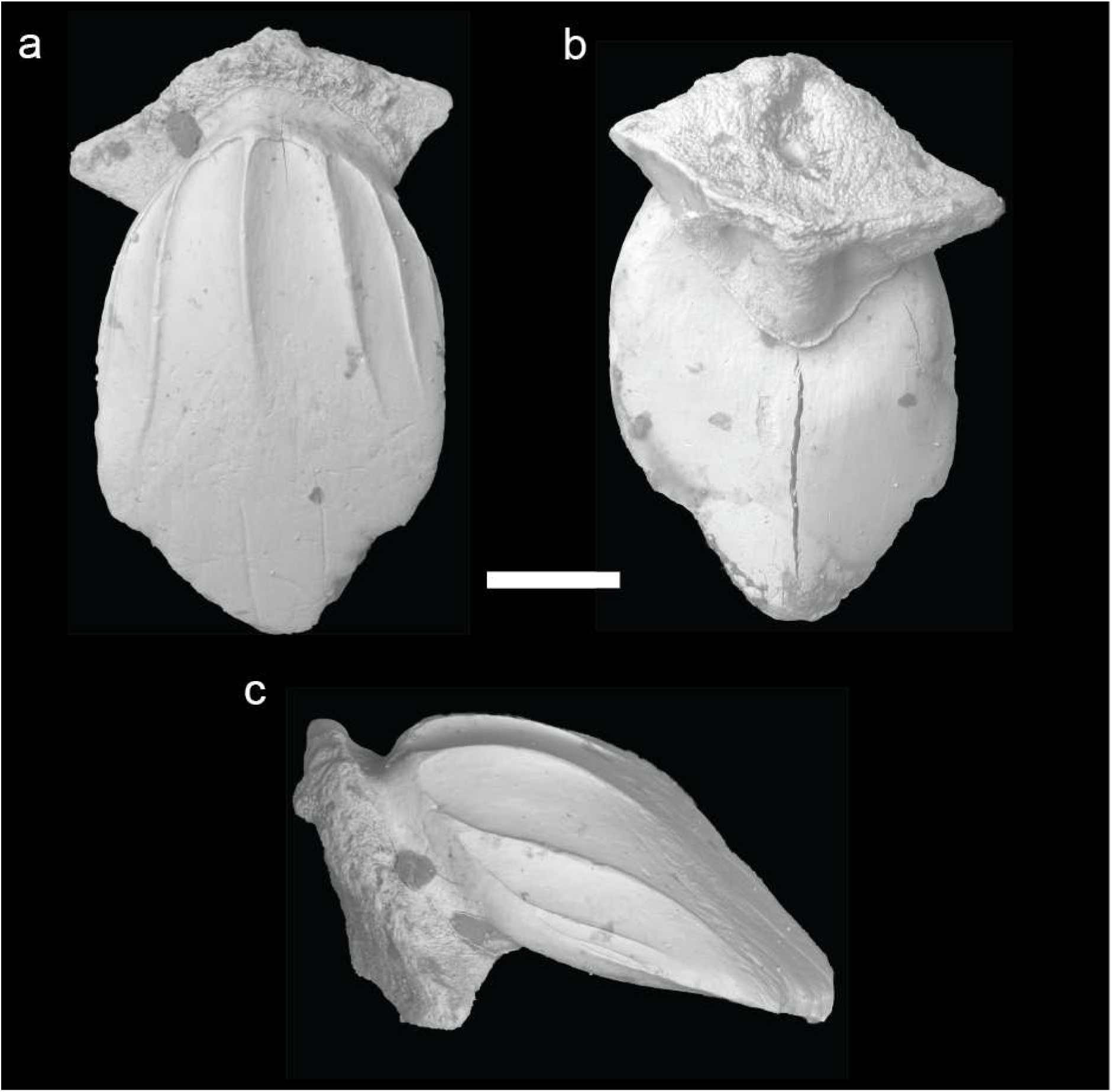
Isolated dermal denticle of *Pararhincodon n. sp.* NHMUK PV P 17223B imaged with SEM: a, dorsal view; b, ventral view; c, left lateral view. Scale bar 0.2 mm.

**Supplementary Figure 7.**
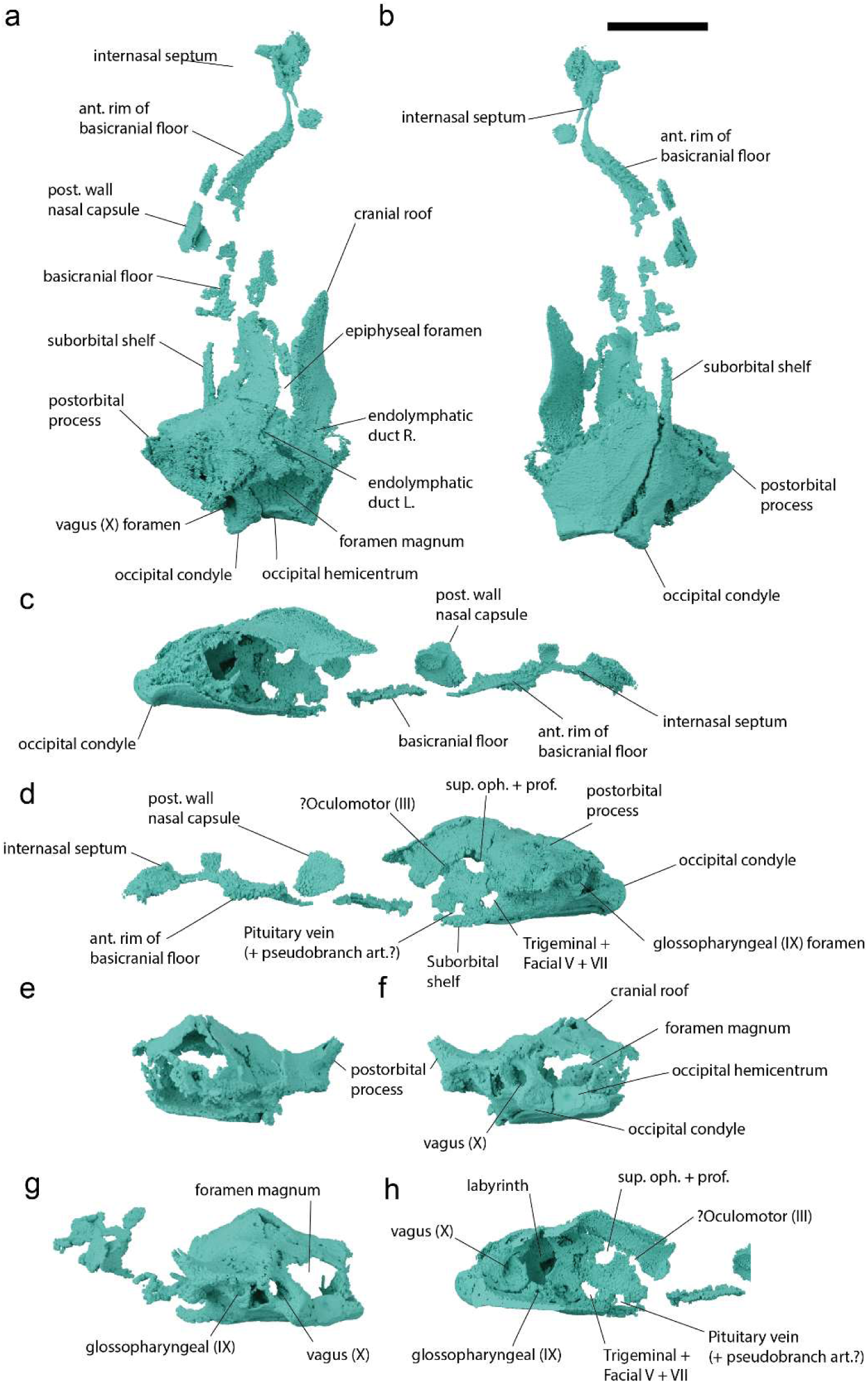
*Pararhincodon n. sp.* NHMUK PV P 17223 neurocranium in a, dorsal; b, ventral; c, left lateral; d, right lateral; e, anterior; f, posterior; g, left posterolateral view; and h, left lateral view bisected to show internal anatomy. All scale bars 10 mm.

**Supplementary Figure 8.**
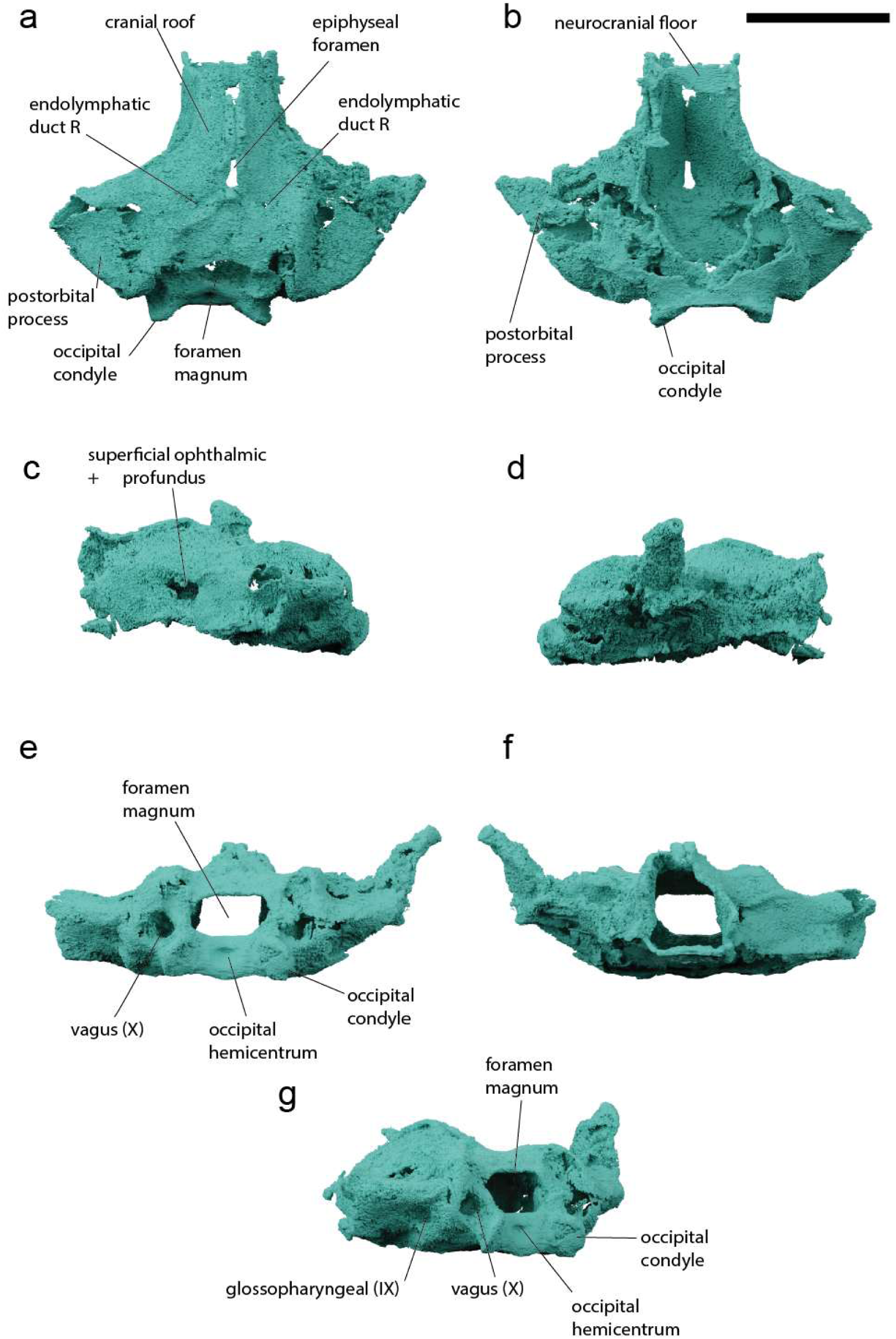
*Pararhincodon n. sp.* NHMUK PV P 73821a neurocranium in a, dorsal; b, ventral; c, left lateral; d, right lateral; e, posterior; f, anterior; and g, left posterolateral views. All scale bars 10 mm.

**Supplementary Figure 9.**
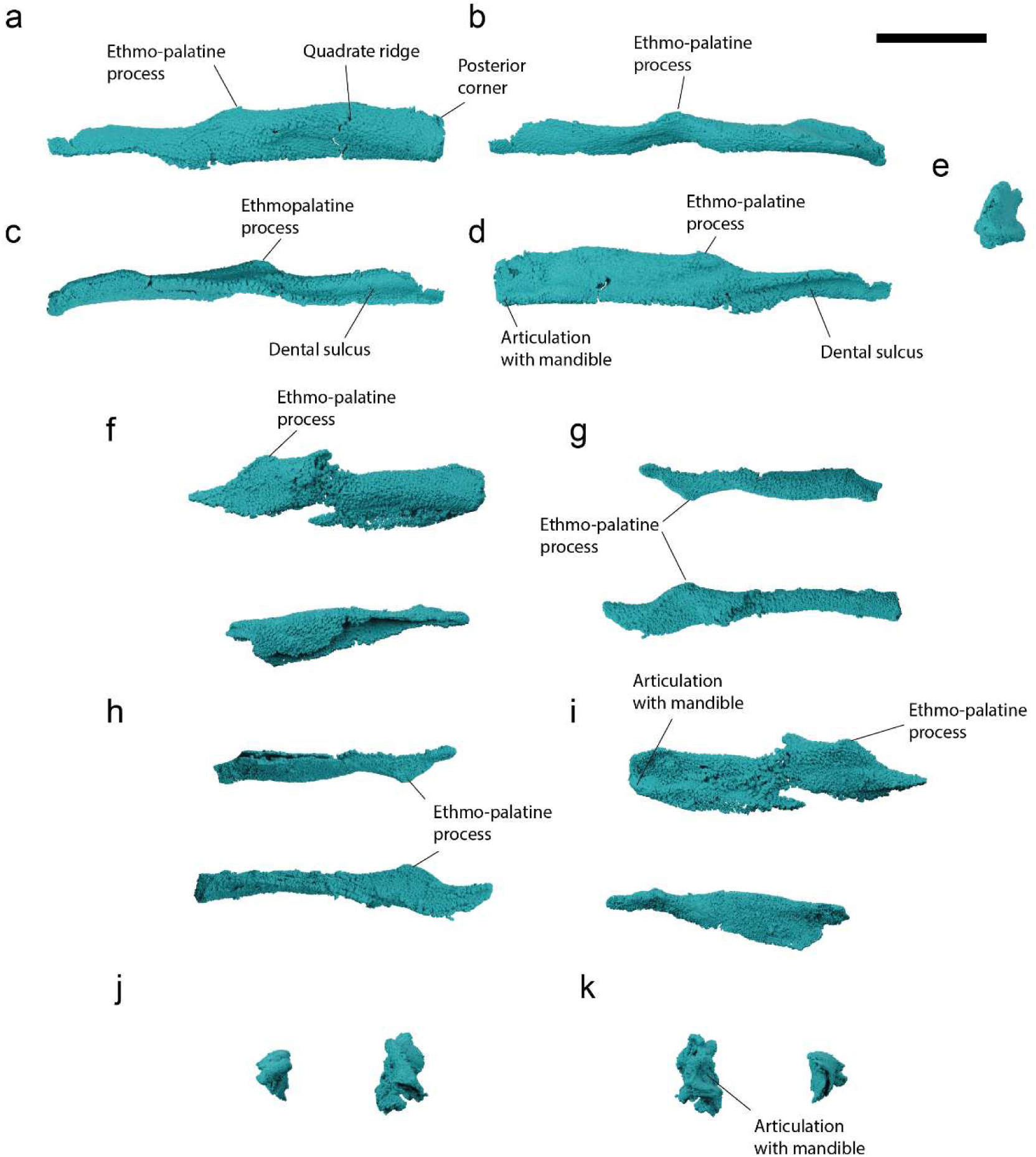
*Pararhincodon n. sp.* palatoquadrate. a-f, left palatoquadrate of *P*. *n. sp.* NHMUK PV P 17223 in labial (a), dorsal (b), ventral (c), lingual (d), and posterior (e) views; f-k, Meckel’s cartilages of *P. n. sp.* NHMUK PV P 73821A in labial (f), dorsal (g), ventral (h), lingual (i), anterior (j) and posterior (k) views. Scale bar 10 mm.

**Supplementary Figure 10.**
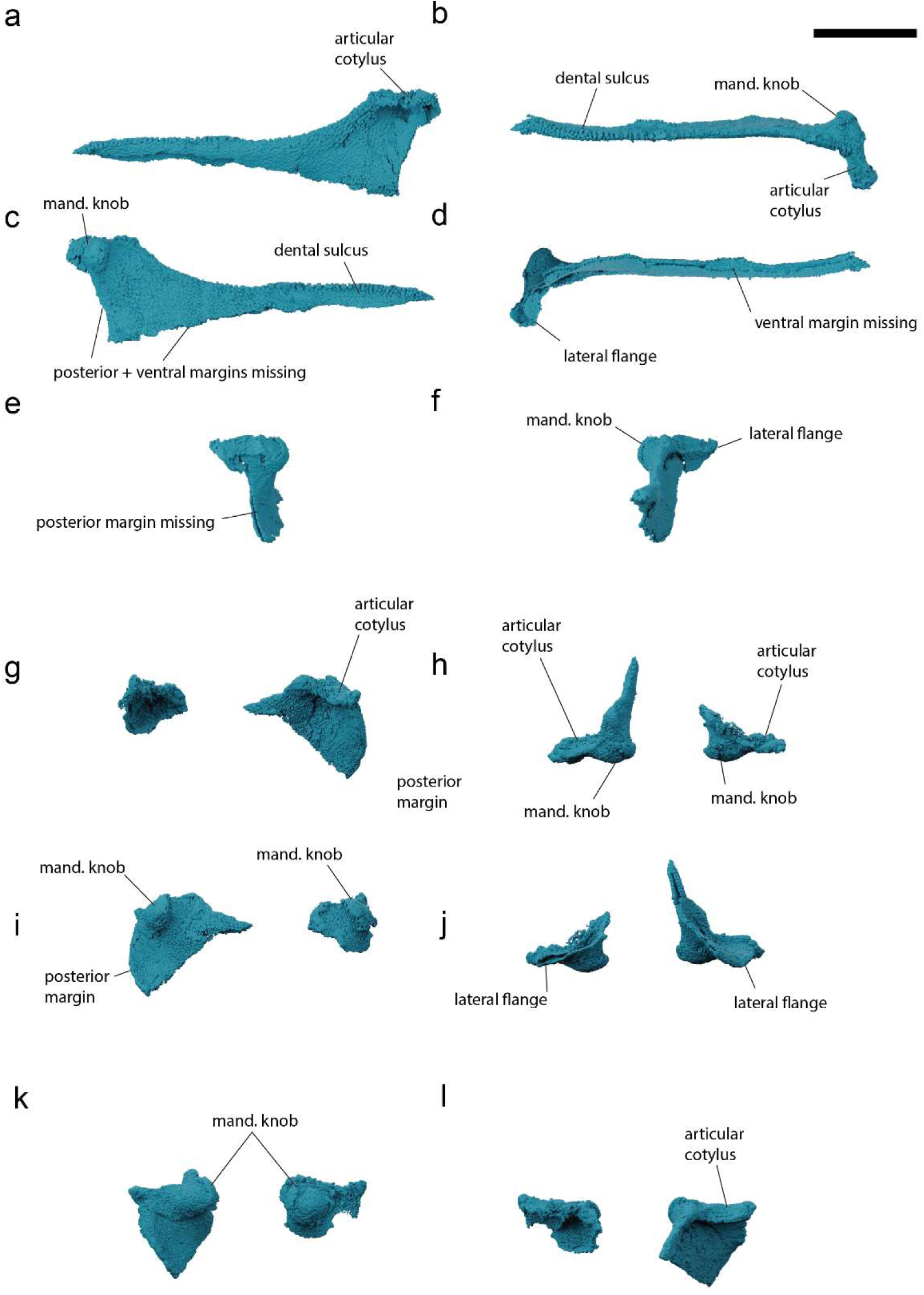
*Pararhincodon n. sp.* Meckel’s cartilage. A-F, left Meckel’s cartilage of *P. n. sp.* NHMUK PV P 17223 in labial (a), dorsal (b), lingual (c), ventral (d), posterior (e), and anterior (f) view; g-l Meckel’s cartilages of *P. n. sp.* NHMUK PV P 73821a in labial (g), dorsal (h), lingual (i), ventral (j), posterior (k), and anterior (l) views. Scale bar 10 mm.

**Supplementary Figure 11.**
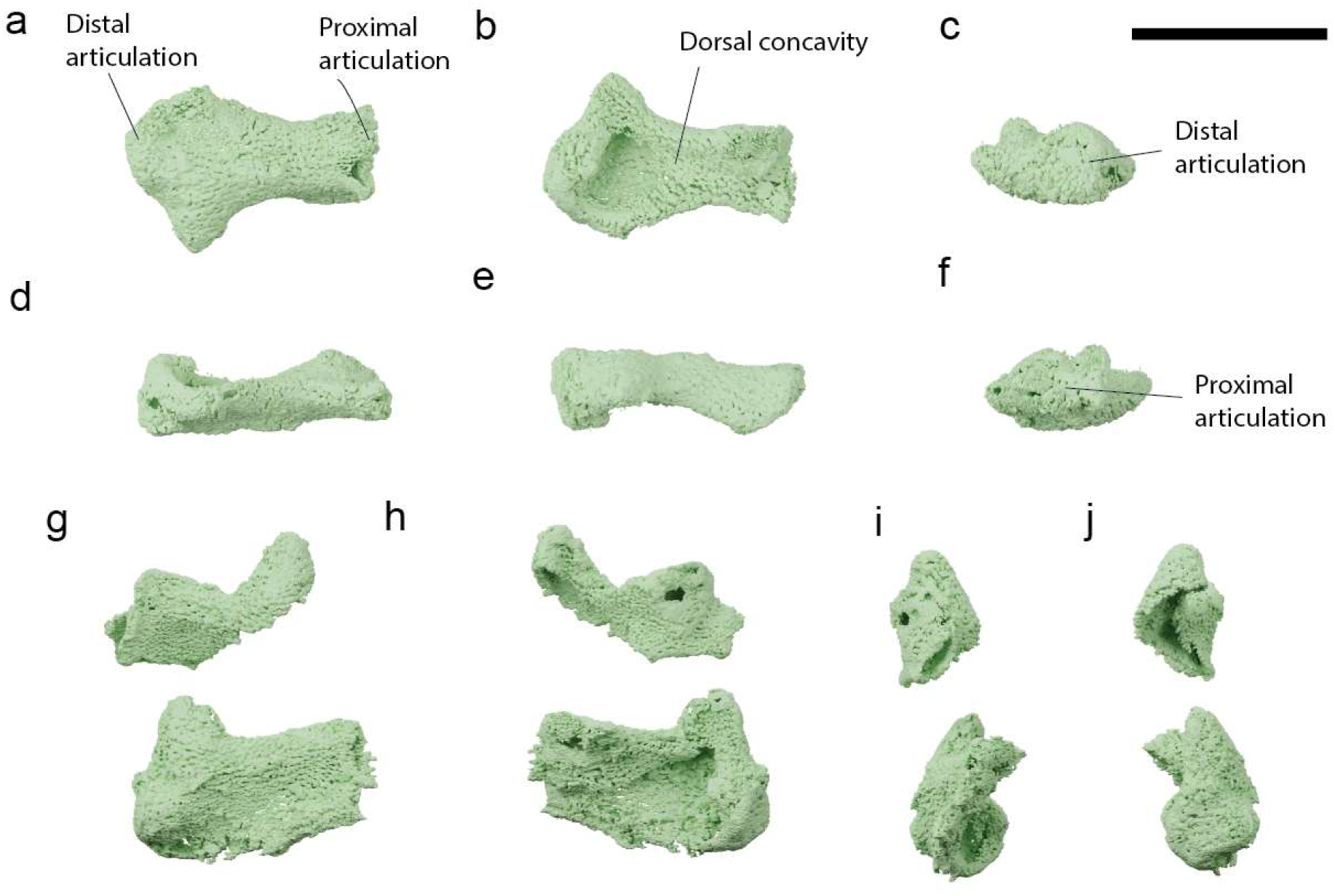
*Pararhincodon n. sp.* hyomandibula. a-f, left hyomandibula of *P*. *n. sp.* NHMUK PV P 17223 in ventral (a), dorsal (b), distal (c), anterior (d), posterior (e), and proximal (f) views; g-j, hyomandibulae of *P. n. sp.* NHMUK PV P 73821a in ventral (e), dorsal (f), proximal (i), and distal (j) views. Scale bar 10 mm.

**Supplementary Figure 12.**
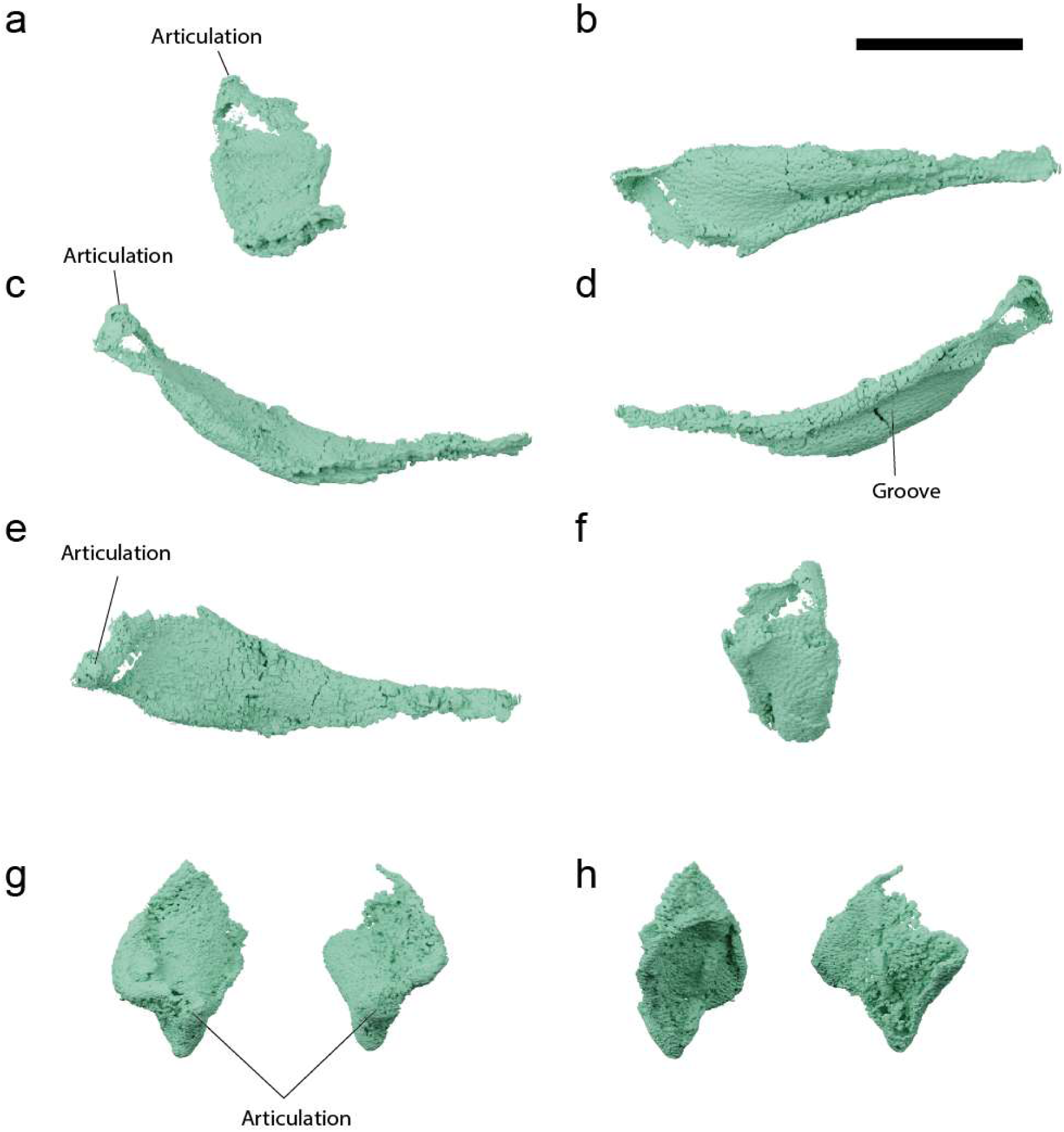
*Pararhincodon n. sp.* ceratohyal. a-f, left ceratohyal of *P. n. sp.* NHMUK PV P 17223 in anterior (a), ventral (b), right lateral (c), left lateral (d), dorsal (e), and posterior (f) views; g-h, ceratohyals of *P. n. sp.* NHMUK PV P 73821a in dorsal (g), and ventral (h) views. Scale bar 10 mm.

**Supplementary Figure 13.**
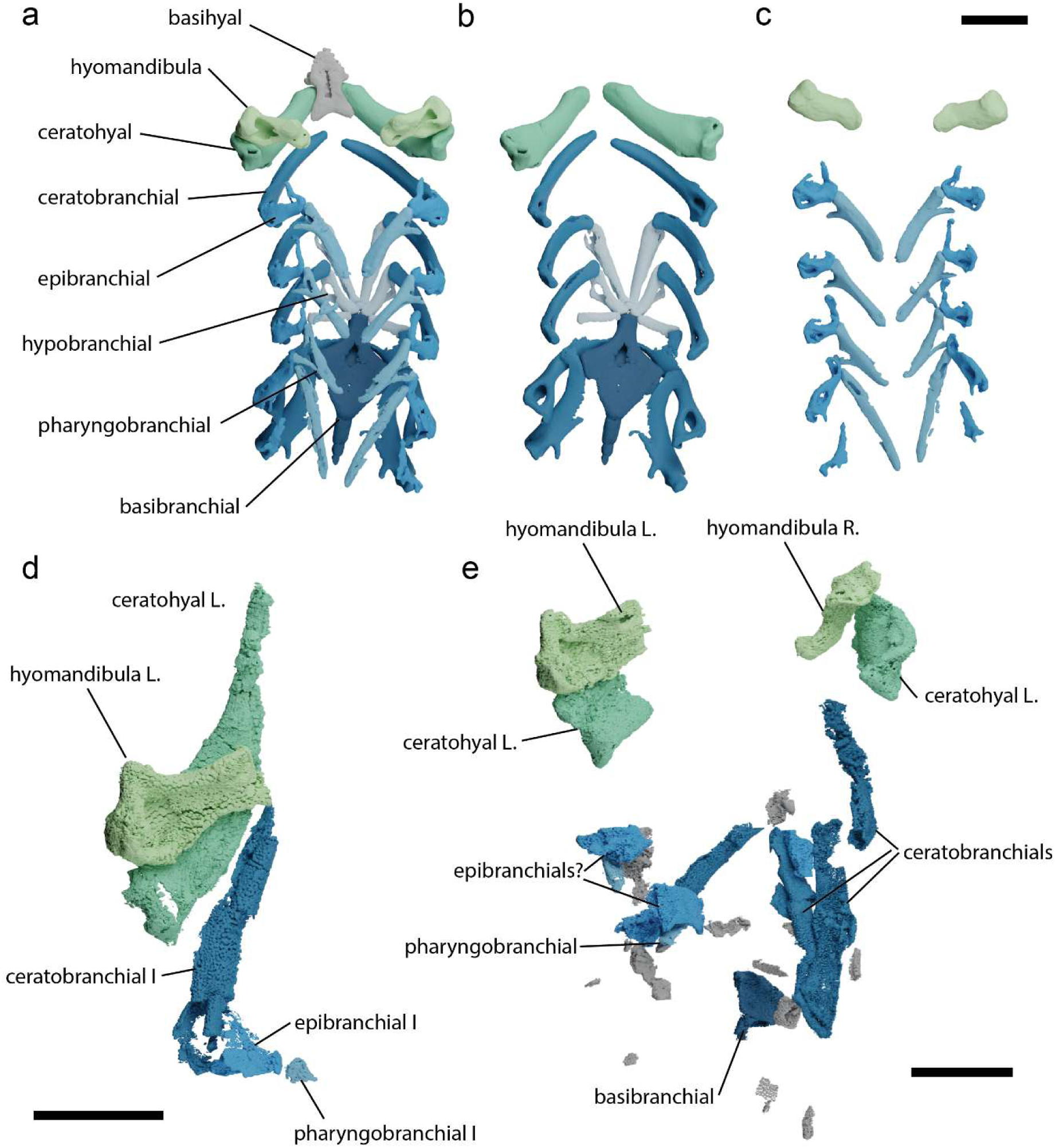
Pharyngeal skeleton of *Pararhincodon n. sp.* NHMUK PV P 73821a by comparison to that of *Parascyllium variolatum CSIRO CA 3311.* a, pharyngeal skeleton of *P. variolatum* in dorsal view; b, ventral components in dorsal view; c, dorsal components in ventral view; d, pharyngeal skeleton of *P. n. sp.* NHMUK PV P 17223 in dorsal view; e, pharyngeal skeleton of *P. n. sp.* NHMUK PV P 73821a in dorsal view, unidentifiable fragments of cartilage in grey. Scale bars = 10 mm

**Supplementary Figure 14.**
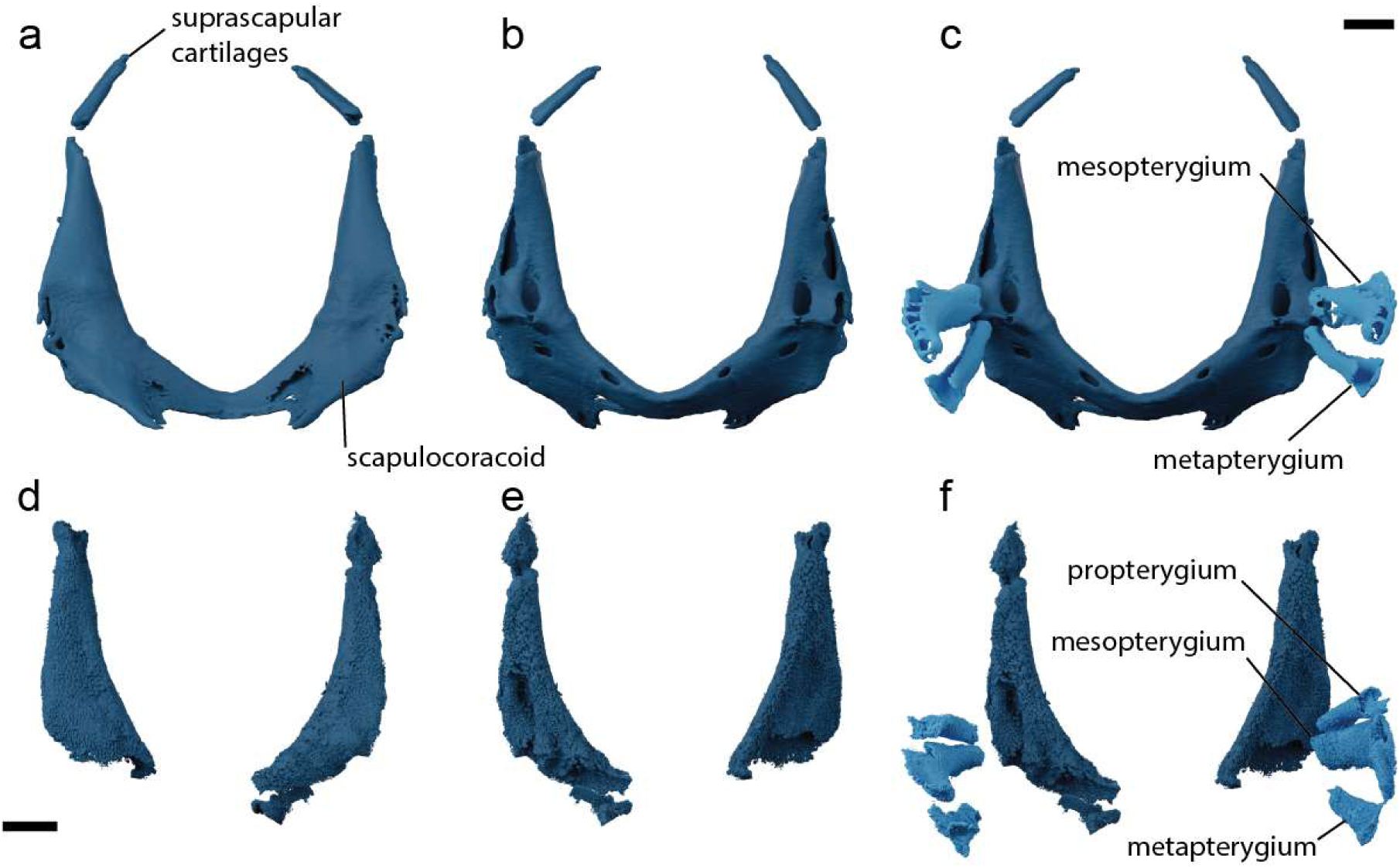
Pectoral skeleton of *Pararhincodon n. sp.* by comparison to that of *Parascyllium variolatum CSIRO CA 3311.* a-b, scapulocoracoid of *Parascyllium variolatum* in anterior (a) and posterior (b) views; c, scapulocoracoid and pectoral basipterygia of *Parascyllium variolatum* in posterior view; d-e main scapulocoracoid chondrifications of *P. n. sp.* NHMUK PV P 73821a in anterior (d) and posterior (e) views; f, scapulocoracoid and pectoral basipterygia of *P. n. sp.* NHMUK PV P 73821A in approximate life position. Scale bars = 5 mm

**Supplementary Figure 15.**
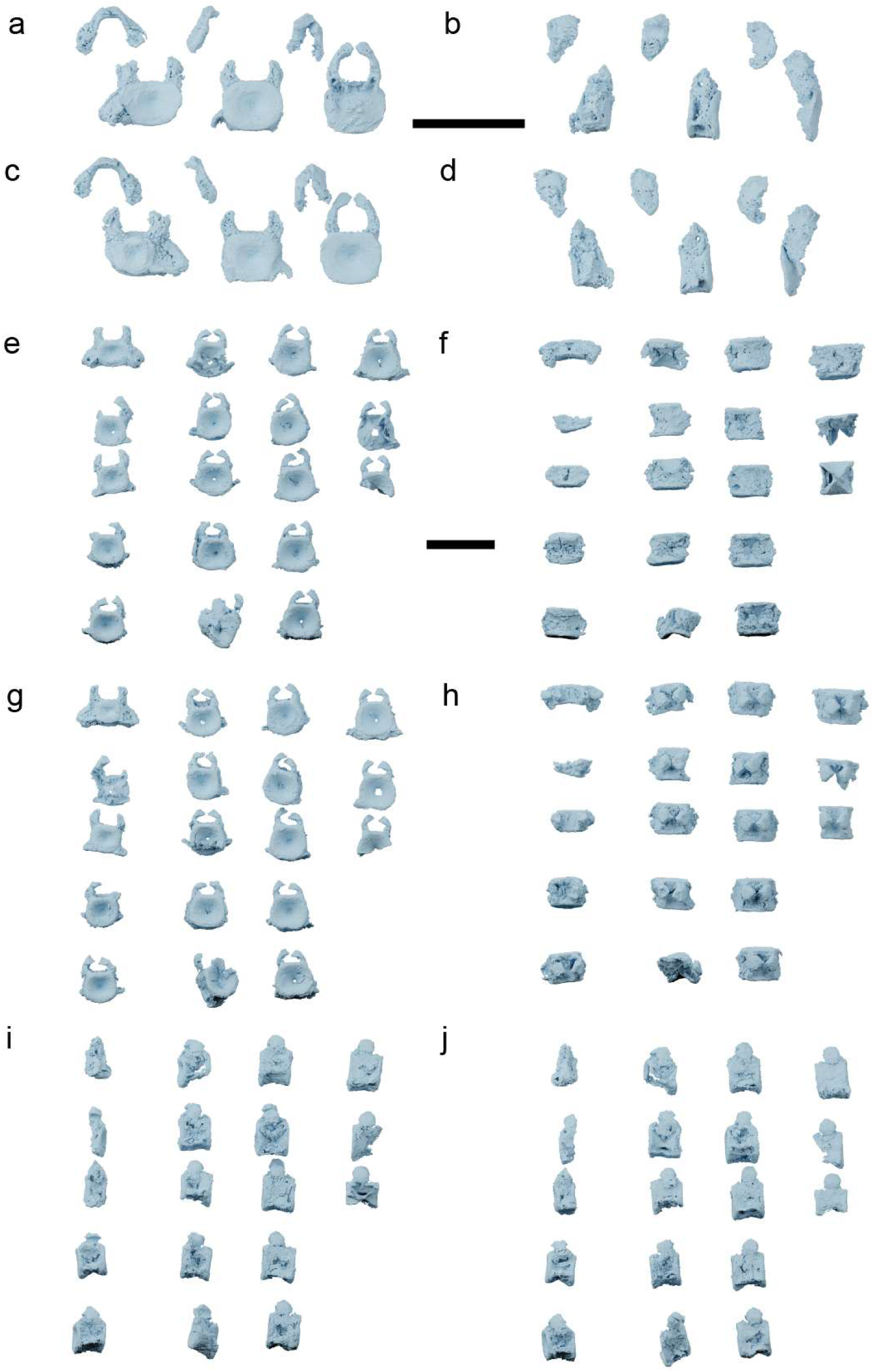
*Pararhincodon n. sp.* Vertebrae. a-d, vertebrae of *P. n. sp.* NHMUK PV P 17223 in anterior (a), right lateral (b), posterior (c), and left lateral (d) views; e- j vertebrae of *P. n. sp.* NHMUK PV P 73821a in anterior (e), ventral (f); posterior (g); dorsal (h) left lateral (i); and right lateral (j) views. All scale bars 10 mm.

